# Mucosal immunoglobulins protect the olfactory organ of teleost fish against parasitic infection

**DOI:** 10.1101/380691

**Authors:** Yong-Yao Yu, Wei-Guang Kong, Ya-Xing Yin, Fen Dong, Zhen-Yu Huang, Guang-Mei Yin, Shuai Dong, Irene Salinas, Yong-An Zhang, Zhen Xu

**Author notes:** These authors contributed equally to this work (Yong-Yao Yu, Wei-Guang Kong).

## Abstract

The olfactory organ of vertebrates receives chemical cues present in the air or water and, at the same time, they are exposed to invading pathogens. Nasal-associated lymphoid tissue (NALT), which serves as a mucosal inductive site for humoral immune responses against antigen stimulation, is present in teleosts and mammals. IgT in teleosts is responsible for similar functions to those carried by IgA in mammals. Moreover, teleost NALT is known to contain B-cells and teleost nasal mucus contains immunoglobulins (Igs). Yet, whether nasal B cells and Igs respond to infection remains unknown. We hypothesized that water-borne parasites can invade the nasal cavity of fish and elicit local specific immune responses. To address this hypothesis, we developed a model of bath infection with the *Ichthyophthirius multifiliis* (Ich) parasite in rainbow trout, *Oncorhynchus mykiss*, an ancient bony fish, and investigated the nasal adaptive immune response against this parasite. Critically, we found that Ich parasites in water could be reach the nasal cavity and successfully invade the nasal mucosa. Moreover, strong parasite-specific IgT responses were exclusively detected in the nasal mucus, and the accumulation of IgT^+^ B-cells was noted in the nasal epidermis after Ich infection. Strikingly, local IgT^+^ B-cell proliferation and parasite-specific IgT generation were found in the trout olfactory organ, providing new evidence that nasal-specific immune responses were induced locally by a parasitic challenge. Overall, our findings suggest that nasal mucosal adaptive immune responses are similar to those reported in other fish mucosal sites and that an antibody system with a dedicated mucosal Ig performs evolutionary conserved functions across vertebrate mucosal surfaces.

**Author Summary:** The olfactory organ is a vitally important chemosensory organ in vertebrates but it is also continuously stimulated by pathogenic microorganisms in the external environment. In mammals and birds, nasopharynx-associated lymphoid tissue (NALT) is considered the first line of immune defense against inhaled antigens and in bony fish, protecting against water-borne infections. However, although B-cells and immunoglobulins (Igs) have been found in teleost NALT, the defensive mechanisms of parasite-specific immune responses after pathogen challenge in the olfactory organ of teleost fish remain poorly understood. Considering that the NALT of all vertebrates has been subjected to similar evolutionary forces, we hypothesize that mucosal Igs play a critical role in the defense of olfactory systems against parasites. To confirm this hypothesis, we show the local proliferation of IgT^+^ B-cells and production of pathogen-specific IgT within the nasal mucosa upon parasite infection, indicating that parasite-specific IgT is the main Ig isotype specialized for nasal-adaptive immune responses. From an evolutionary perspective, our findings contribute to expanding our view of nasal immune systems and determining the fate of the host–pathogen interaction.

## Introduction

Olfaction is a vital sense for all animals [1]. To receive an olfactory signal, terrestrial vertebrates inhale gases containing volatile chemical substances, while aquatic vertebrates like teleost fish actively draw water containing dissolved chemicals into the olfactory organs [2]. Simultaneously, during this process, the olfactory organs are constantly stimulated by toxins and pathogens in the air or water [3]. Therefore, there is an evident need to defend the large, delicate surface of olfactory organs from pathogenic invasion.

In mammals, nasopharynx-associated lymphoid tissue (NALT), is a paired mucosal lymphoid organ containing well-organized lymphoid structures (organized MALT, O-MALT) and scattered or disseminated lymphoid cells (diffuse MALT, D-MALT) and is traditionally considered the first line of defense against external threats [4]. Similar to the Peyer’s patches in the guts of mammals, O-NALT has distinct B-cell zones [5, 6], and humoral immune responses occur in response to infection or antigenic stimulation [7]. Importantly, the higher percentage of IgA^+^ B-cells in D-NALT compared with that in O-NALT indicates that D-NALT may play an important role in nasal antibody-mediated immunity [8]. Interestingly, NALT in early vertebrates like teleost fish has structures and components similar to those of mammalian NALT [1]. Teleost NALT has thus far been described as D-NALT but lacks O-NALT. Teleost NALT includes B-cells, T cells, myeloid cells and expresses innate and adaptive related molecules [9]. Thus, from an evolutionary viewpoint, NALT in teleost fish is equipped to rapidly respond to antigens present in the water environment [3].

Teleost fish represent the most ancient bony vertebrates with a nasal-associated immune system [10] and containing immunoglobulins (Igs) [9]. So far, only three Ig classes (IgM, IgD, and IgT/Z) have been identified in teleosts [11]. Teleost IgM has been considered the principal Ig in plasma, and strong parasite-specific IgM responses have been induced in systemic immunity [12–14]. Although secreted IgD (sIgD) has been found in the coating of a small percentage of the microbiota at the gill mucosa surface, its function remains unknown [15]. In contrast, teleost IgT (also called IgZ in some species) has been identified at the genome level and found to play a specialized role in response to pathogen infection in mucosal tissues [15–17]. Moreover, IgT^+^ B-cells represent the predominant mucosal B-cell subset, and the accumulation of IgT^+^ B-cells has been detected after infection in trout gut-, skin-, and gill-associated lymphoid tissues (GALT, SALT, and GIALT) [15–17]. Interestingly, in mammals, parasite-specific IgA has been mainly induced after pathogenic infection, and it has mediated nasal-adaptive immunity [18–20]. However, in teleosts, the role of the three Ig classes and B-cells in the olfactory organ is still unknown. Thus, given the abundance of IgT^+^ B-cells as well as the high concentration of IgT in the olfactory organ [9, 21], we hypothesized that IgT is the major Ig involved in the pathogen-specific immune responses in the NALT of teleost fish.

To test the aforementioned hypothesis, here, we studied the nasal B-cell and parasite-specific Ig responses to the ciliated parasite *Ichthyophthirius multifiliis* (Ich) in rainbow trout, a model species in the field of evolutionary and comparative immunology [22, 23]. Our findings show that the olfactory system of rainbow trout is an ancient mucosal surface that elicits strong innate and adaptive immune responses to Ich infection. In addition, we demonstrate that IgT is the main Ig isotype playing a critical role in nasal adaptive immune responses. Furthermore, we show for the first time the local production of IgT at the nasal mucosa and proliferation of IgT^+^ B-cells after a parasitic challenge in the olfactory organ of teleost fish. These results demonstrate that NALT is both an inductive and effector immune site in teleost fish.

## Results

### Igs in olfactory organ of rainbow trout

Here, nasal mucosa IgT was detected by Western blot consistent with the reported molecular mass using anti-trout IgT antibody [15–17]. To understand the protein characterization of nasal IgT, we collected the nasal mucosa of rainbow trout and loaded 0.5 μl of processed mucus into a gel filtration column. From these results, we found that a portion of IgT in the nasal mucosa was present in polymeric form, as it eluted at a fraction similar to that of trout nasal IgM, a tetrameric Ig (Fig 1A), and simultaneously, some IgT was consistently eluted in monomeric form. Next, nasal mucosa polymeric IgT (pIgT) migrated to the same position as a monomer by SDS-PAGE under non-reducing conditions, indicating that nasal pIgT is associated by non-covalent interactions (Fig 1B, right panel). However, unlike IgM and IgT, IgD in nasal mucosa eluted at 8.5 and 9.5 as a monomer at the molecular weight range previously studied for serum IgD [15]. Using Western blot, under the same immunoblot conditions, we found that nasal IgD and IgM migrated as a monomer and polymer, respectively (Fig 1B, left and middle panel). These findings are similar to those previously reported in the gut [16], skin [17], and gill [15]. Finally, using Western blot, we compared and analyzed the concentrations of three Igs and the ratios of IgT/IgM and IgD/IgM in nasal mucus and serum (Fig 1C–F), respectively. Our results showed that the protein concentration of IgT was ~ 164- and ~ 602-fold lower than that of IgM in nasal mucus and serum, respectively (Fig 1C and D). Although IgM was found to be the highest Ig in nasal mucosa, the IgT/IgM ratio in nasal mucus was ~ 4-fold higher than that in serum (Fig 1E), whereas there was no obvious difference between them in terms of the IgD/IgM ratio (Fig 1F).

**Fig 1.**
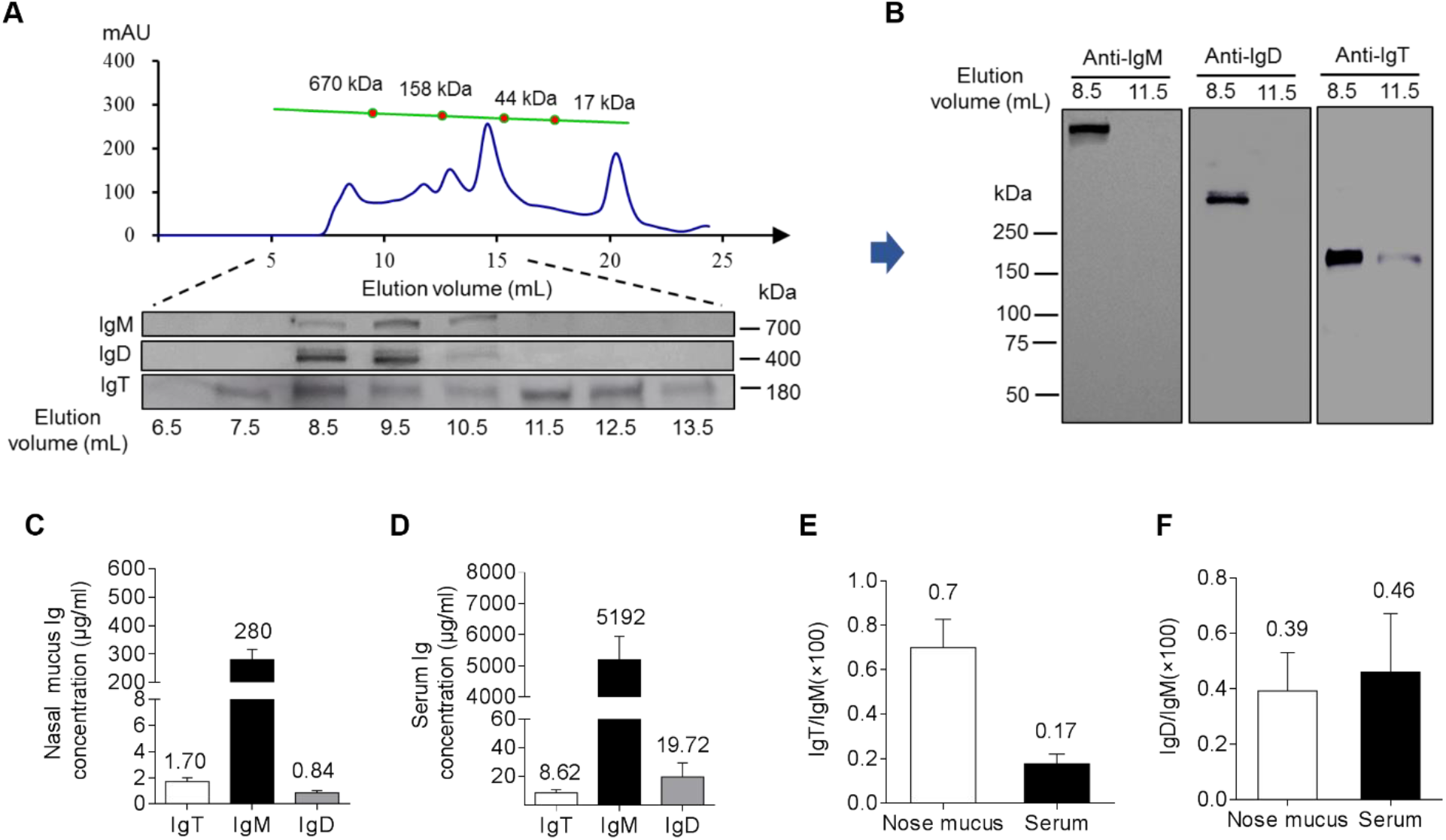
Structural characterization of immunoglobulins in trout nasal mucus. (A) Fractionation of nasal mucus (~ 0.5 ml) by gel filtration (upper) followed by immunoblot analysis of the fractions with anti-trout IgM-, anti-trout IgD-specific mAbs, and anti-trout IgT-specific pAbs (lower). *A_280_*, absorbance at 280 nm. (B) SDS-PAGE of gel-filtration fractions (4-15 %) corresponding to elution volumes of 8.5 ml and 11.5 ml under non-reducing conditions followed by immunoblot analysis with anti-trout IgM-, anti-trout IgD-specific mAbs or anti-trout IgT-specific pAbs. Immunoblot and densitometric analysis of the concentration of IgT, IgM and IgD in nasal mucus (C) and serum (D) (*n* = 12 fish). Ratio of IgT to IgM concentration (E) and IgD to IgM concentration (F) in nasal mucus and serum, calculated from the values shown in C and D. Results in Fig C-F are expressed as mean and s.e.m. obtained from 12 individual fishes.

### Polymeric Ig receptor (pIgR) in olfactory organ of rainbow trout

In mammals, pIgR can mediate the transepithelial transport of secretory IgA (sIgA) into the nasal mucosa [27, 28]. In trout, we previously found that the secretory component of trout pIgR (tSC) is associated with secretory IgT (sIgT) in the gut [16], skin [17], and gills [15] and pIgR expression is very high in control rainbow trout olfactory organ [9]. Here, using pIgR polyclonal antibody [16], tSC was detected in the nasal mucosa but not in the serum (Fig 2A). By coimmunoprecipitation assay and immunoglobulins, we showed that antibodies against trout IgT was able to coimmunoprecipitate tSC in nasal mucus (Fig 2B). Moreover, using immunofluorescence microscopy, most of the pIgR-containing cells were located in the OE of trout, some of which were stained with IgT (Fig 2C; isotype-matched control antibodies (S1A Fig).

**Fig 2.**
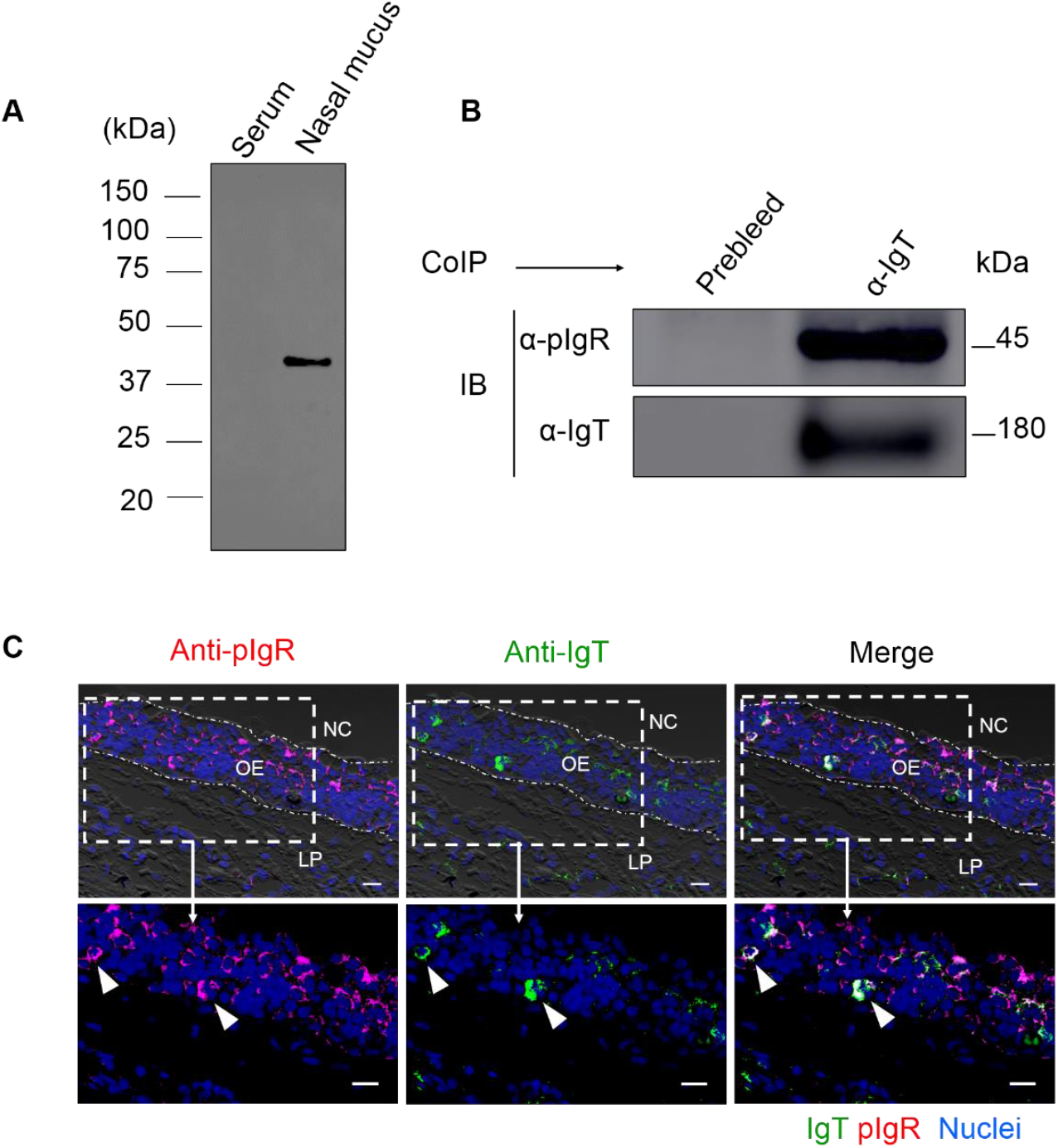
Trout pIgR associates nasal sIgT. (A) SDS-PAGE under reducing conditions of trout serum and nasal mucus (~ 5 μg each), followed by immunoblot analysis using anti-trout pIgR antibody. (B) Co-immunoprecipitation (CoIP) of nasal mucus with anti-trout IgT antibody, followed by immunoblot analysis under reducing conditions (pIgR detection, upper panels) or non-reducing conditions (IgT detection, lower panels). (C) Immunofluorescence staining for pIgR with IgT in olfactory organ paraffinic sections of rainbow trout. Differential interference contrast images of olfactory organ paraffin sections were stained with anti-trout pIgR (magenta), anti-trout IgT (green) and DAPI for nuclei (blue) (*n* = 9) (isotype-matched control antibodies for anti-pIgR in S1A Fig). Enlarged sections of the areas outlined showing some pIgR/IgT colocalization (white arrowhead). NC, nasal cavity; OE, olfactory epithelium; LP, lamina propria. Scale bar, 20 μm. Data are representative of at least three different independent experiments.

### Response of immune-related genes in trout olfactory organ to Ich parasite infection

To evaluate whether the trout olfactory organ expresses immune-related genes after pathogen challenge, we firstly selected the Ich parasite bath infection model and investigated nasal mucosal immune responses (S2A Fig). Ich is a parasite that directly invades the mucosal tissues of fish, such as the skin, gill, and fin, and it might elicit a strong immune response [15, 17], however, it has never been reported to infect the olfactory organ of fish. At 7 days post-infection, the phenotype of the small white dots appeared on the trout’s skin and fin surface (S2B Fig), and by examining paraffin sections of olfactory organs stained with H & E, the Ich parasite was found within the nasal cavity and mucosa, interestingly, most of which were present in lateral regions compared with tips of nasal lamina propria (S2C Fig). In addition, by reverse transcription quantitative real-time PCR (RT-qPCR), we detected the expression of Ich-18SrRNA in the olfactory organ, gills, skin, head kidney, and spleen of trout after 7 days infection and controls (S2D Fig). Ich-18SrDNA expression levels were comparable in the nose and the gills, one of the target organs of Ich, highlighting the importance of Ich nasal infections in trout. A time series study of Ich-18SrRNA expression showed that parasites levels in the nose peaked at day 7 post-infection with a second wave occurring at day 21. Interestingly, Ich levels dropped dramatically on day 28 but increased again 75 days post-infection (S2E Fig). Using RT-qPCR, we measured the expression of 26 immune-related genes in the olfactory organ of trout at days 1, 7, 14, 21, 28, and 75 post-infection. Overall, greatest changes in expression of pro-inflammatory and complement-related genes as well as, occurred at 7 days post-infection (Fig 3A, primers used are shown in S1 Table) when parasite levels were highest. Expression of IgT and IgM heavy chain genes, in turn, increased later during infection, starting at day 21 and peaking at day 28, whereas no obvious change in IgD heavy chain expression was observed (Fig 3A and B). IgT expression was the most up-regulated (~ 258-fold) compared to IgM (~ 116-fold) on day 28 and remained up-regulated on day 75 (~ 112-fold) compared to IgM which was only moderately higher than controls (~ 8-fold) (Fig 3B). Moreover, using histological examination, at 7 days post-infection, lamina propria (LP) of trout in the tip (~ 100 μm) showed a significant enlargement (Fig 3C and D), increased goblet cells (Fig 3C and E) in nasal lamella compared with the control. By days 28 and 75, the tissue reaction was smaller, the LP showed some enlargement and abundant goblet cells appeared in nasal lamella (Fig 3C-E). Combined, these results demonstrate that apart from infecting gills, and skin, Ich is able to chronically infect the trout olfactory organ and induce strong long-lasting IgT responses.

**Fig 3.**
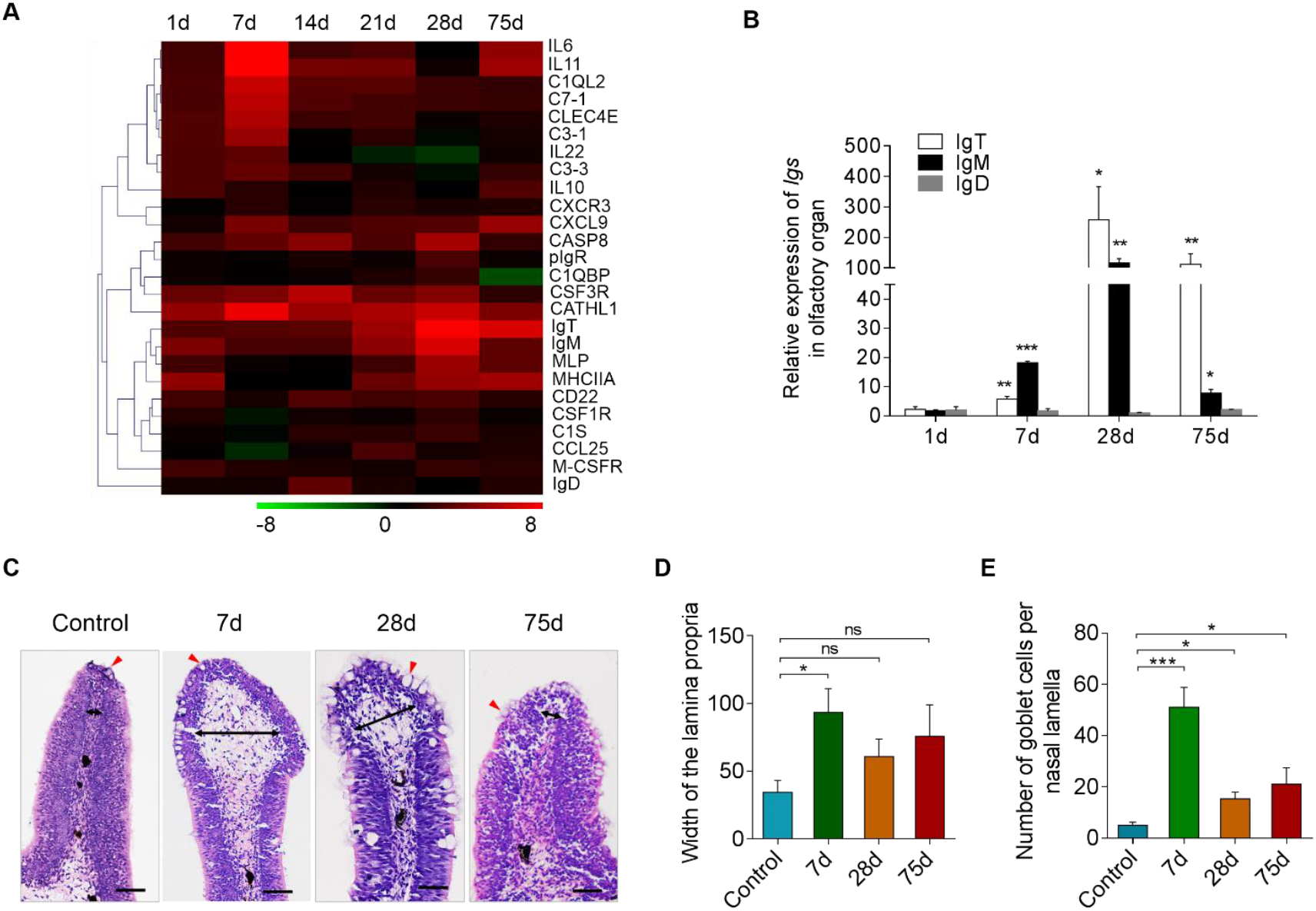
Kinetics of immune response and pathological changes in trout olfactory organ following Ich parasites infection. (A) Heat map illustrates results from quantitative real-time PCR of mRNAs for selected immune markers in parasite-challenged versus control fish measured at 1, 7, 14, 21, 28 and 75 days post infections with Ich parasite in the olfactory organ of rainbow trout (n = 6 fish per group). Color value: log2 (fold change). (B) Relative expression of IgM, IgD and IgT at 1, 7, 21, 28 and 75 days post infection with Ich parasite in olfactory organ of rainbow trout (*n* = 6 fish per group). (C and D) Histological examination (haematoxylin & eosin stain; H & E) (C) of the olfactory organ and the width of the olfactory lamella (D) from ich-infected rainbow trout 7, 21, 28, 75 d.p.i and uninfected fish (n = 6 fish per group). Black arrows indicate the width of LP at the tip (100 μm from the lamellar tip) and medial (250 μm from the lamellar tip) regions of the olfactory lammella and red arrows indicate goblet cells. \**P* < 0.05, \*\**P* < 0.01 and \*\*\**P* < 0.001 (one – way ANOVA with Bonferroni correction). Data are representative of at least three independent experiments (mean and s.e.m.). Anova, analysis of variance.

### Response of Igs in trout olfactory organ to Ich parasite infection

The high expression of IgT in the olfactory organ of trout after an Ich parasite challenge led us to hypothesize a critical role of IgT in nasal immunity. Using immunofluorescent micrographs, Ich trophonts could be easily detected in the olfactory organ of trout after 28 days of infection (Fig 4) using an anti-Ich antibody (isotype-matched control antibodies, S3 Fig). Interestingly, most parasites detected in the olfactory organ of trout were intensely coated with IgT, while only some parasites were slightly coated with IgM and nearly no parasites were coated with IgD (Fig 4). In addition, we found few IgT^+^ and IgM^+^ B-cells in the nasal epithelium of control fish (Fig 5A; isotype-matched control antibodies, S1B Fig left). Interestingly, a moderate increase of IgT^+^ B-cells was observed in the nasal epithelium of trout after 28 days of infection (Fig 5B; isotype-matched control antibodies, S1B middle Fig). It is worth mentioning that a notable accumulation of IgT^+^ B-cells was detected in the nasal epithelium of survivor fish (75 days post-infection) compared with control trout (Fig 5C; isotype-matched control antibodies, S1B right Fig). More importantly, we observed that some IgT^+^ B-cells appeared to be secreting IgT (Fig 5C, white arrows). In contrast, the abundance of IgM^+^ B-cells did not change in the infected and survivor fish when compared to the controls (Fig 5A–C). Next, we analyzed the percentages of IgT^+^ and IgM^+^ B-cells in the olfactory organs of control, infected, and survivor fish. We observed that, similar to the result obtained by immunofluorescence microscopy, the percentages of IgT^+^ B-cells in the infected group (~ 3.66 ± 0.2%) and survivor group (~ 4.43 ± 0.28%) increased significantly compared to those of the control group (~ 1.72 ± 0.08%) (Fig 6A). In contrast, the percentages of nasal IgM^+^ B-cells did not change in the three groups (Fig 6A). Unlike the results in the olfactory organ, the percentage of IgM^+^ B-cells of the head kidney in the infected group (~ 11.34 ± 0.39%) showed a significant increase compared to that of the control group (~ 6.53 ± 0.27%), while the percentage of IgM^+^ B-cells in the survivor group (~ 8.65 ± 0.67%) showed no significant change (Fig 6B). In contrast, the percentages of IgT^+^ B-cells remained unchanged in both the infected groups and the survivor groups (Fig 5B). In agreement with the increased IgT^+^ B-cells observed in the olfactory organ of infected and survivor fish, the IgT concentration in the nasal mucosa of these fish increased by ~ 2- and ~ 6-fold when compared with control fish, respectively. However, IgM and IgD protein concentrations did not change in any fish groups (Fig 6C). In serum, a ~ 5-fold increase of IgT concentration was observed only in the survivor group, whereas that of IgM in both the infected and survivor group increased by ~ 5-fold with respect to control fish (Fig 6D). As expected, the IgD protein concentration did not change significantly in infected or survivor fish (Fig 6D).

**Fig 4.**
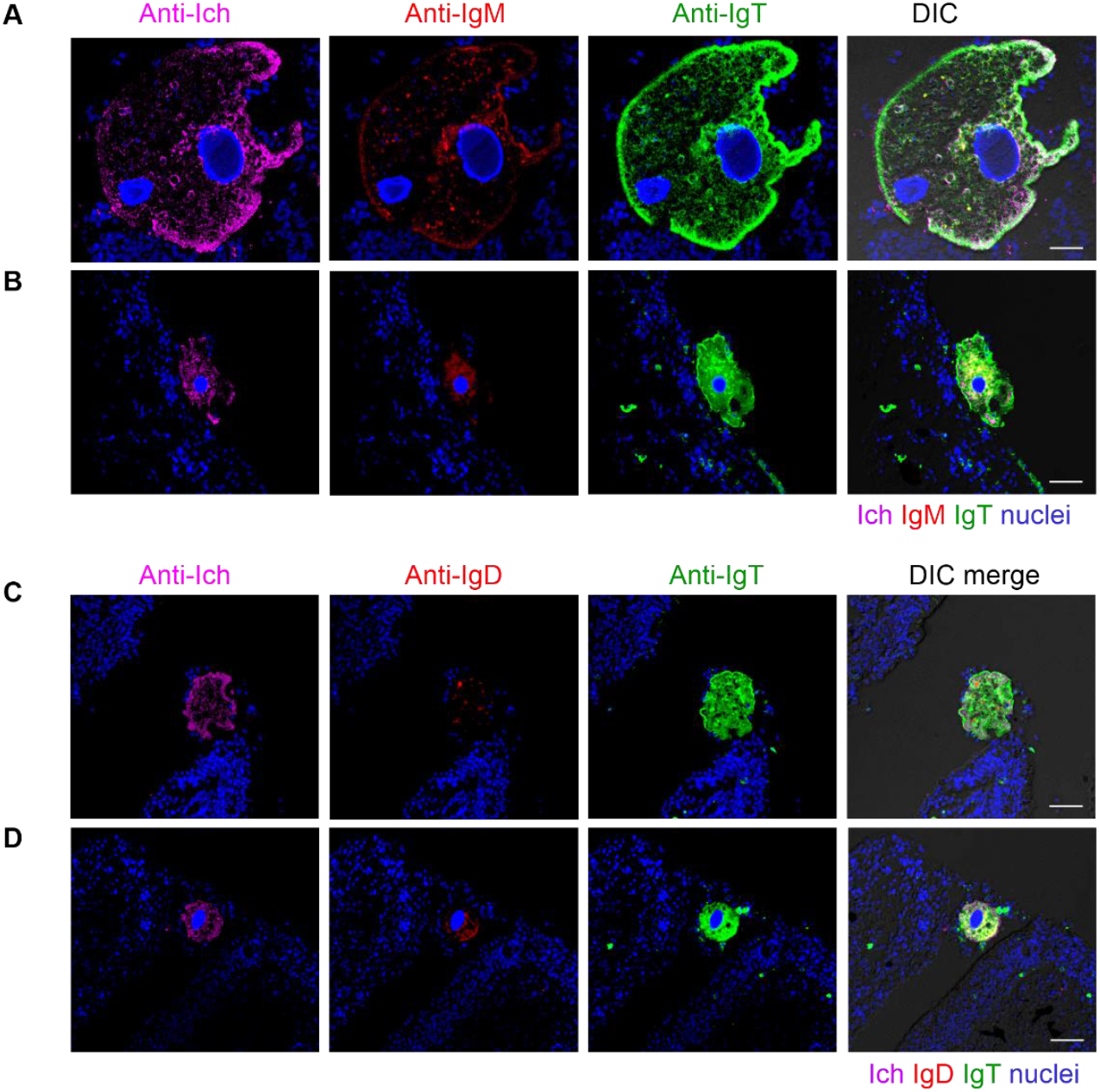
IgT coats Ich parasite located in olfactory organ of infected trout. Four different microscope images (A-D) of slides immunofluorescence staining of Ich parasites in olfactory organ paraffinic sections from trout infected with Ich after 28 days (*n* = 6). (A and B) Immunofluorescence stained with Ich (magenta), IgM (red) and IgT (green), nuclei stained with DAPI (blue) (from left to right). (C and D) Immunofluorescence stained with Ich (magenta), IgD (red) and IgT (green) with nuclei stained with DAPI (blue) (from left to right); DIC images showing merged staining (isotype-matched control antibody staining, S3A-C Fig). Scale bars, 20 μm. Data are representative of at least three different independent experiments.

**Fig 5.**
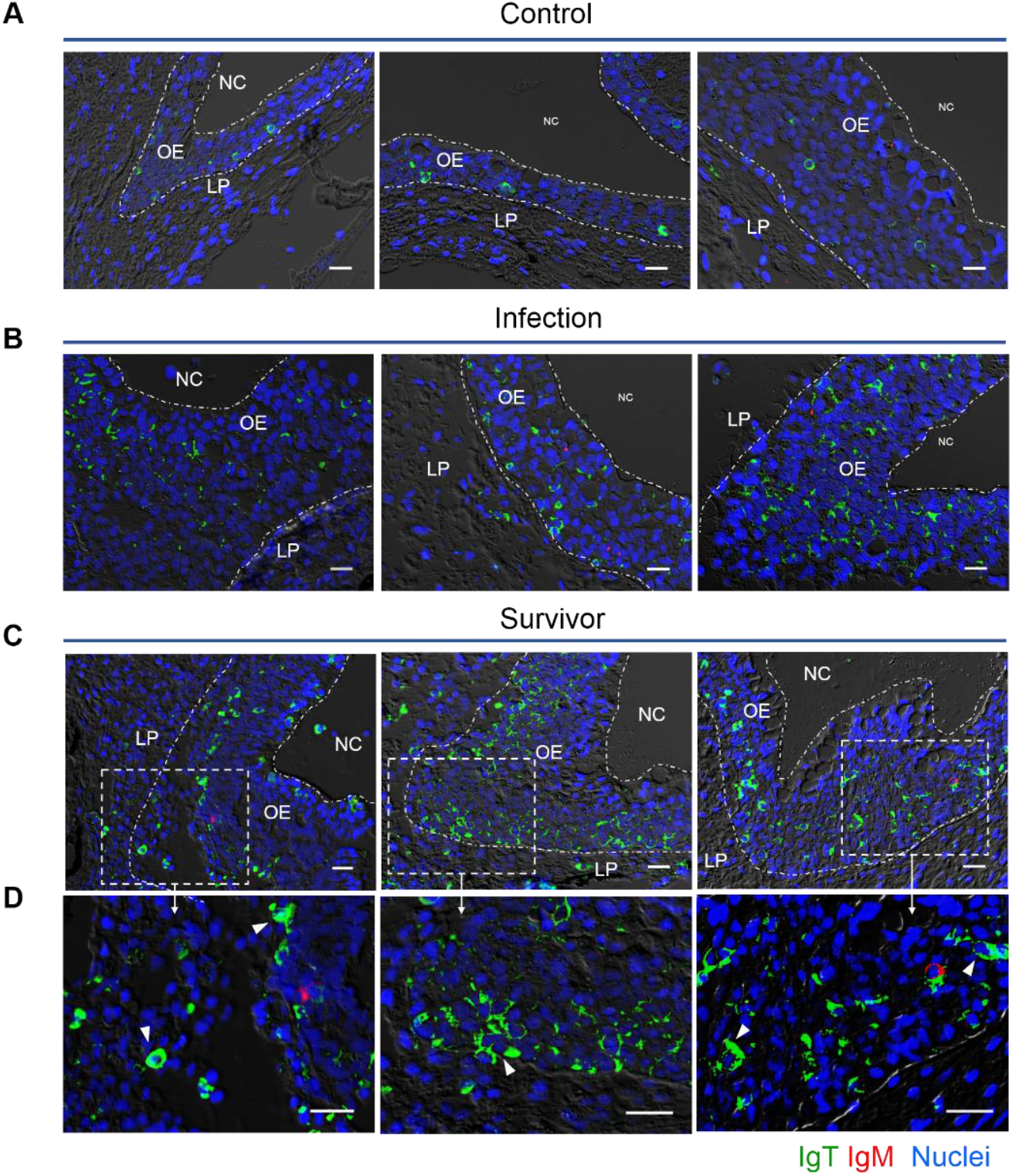
Accumulation of IgT^+^ B cells in the olfactory organ of trout infected with Ich. DIC images of immunofluorescence staining on trout nasal paraffinic sections from uninfected fish (A), 28 days infected fish (B) and survivor fish (C), stained for IgT (green) and IgM (red); nuclei are stained with DAPI (blue). (D) Enlarged images of the areas outlined in c are showing some IgT^+^ B cells possibly secreting IgT (white arrowhead) (isotype-matched control antibody staining, S1B Fig). NC, nasal cavity; OE, olfactory epithelium; LP, lamina propria. Scale bar, 20 μm. Data are representative of at least three different independent experiments (*n* = 8 per group).

**Fig 6.**
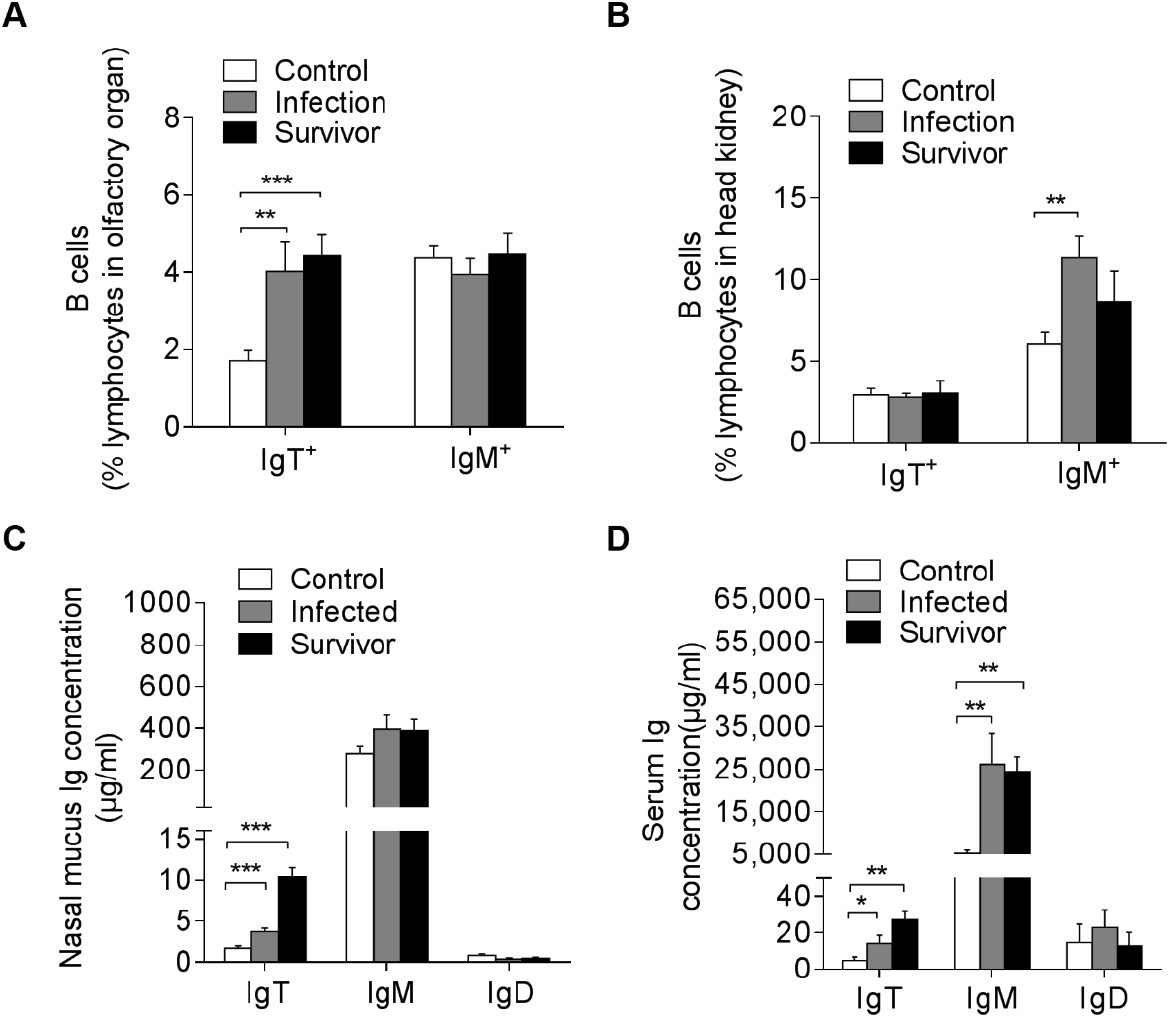
Increases of IgT^+^ B cells and IgT concentration in the olfactory organ of trout infected with Ich. Percentage of IgT^+^ and IgM^+^ B cells in NALT (A) and head kidney (B) leukocytes of uninfected control fish, infected fish and survivor fish measured by flow cytometric analysis (*n* = 12 per group). Concentration of IgT, IgM and IgD in nasal mucus (C) and serum (D) of control, infected and survivor fish (n = 12 per group). \**P* < 0.05, \*\**P* < 0.01 and \*\*\**P* < 0.001 (one - way ANOVA with Bonferroni correction). Data are representative of at least three independent experiments (mean and s.e.m.). Anova, analysis of variance.

The results of large increases of IgT^+^ B-cells and IgT protein levels in the olfactory organs of infected and survivor fish, together with the observation that parasites in the olfactory organ of infected fish appear intensely coated with IgT, suggested that parasite-specific IgT might be secreted in the nasal mucosa response to Ich infection. To verify this hypothesis, using a pull-down assay, we measured the capacity of nasal Igs to bind the Ich parasite (Fig 7). We found a significant increase in parasite-specific IgT binding in up to 1/10 (~ 3.2-fold) and 1/100 (~ 2.8-fold) of the diluted nasal mucus of infected (Fig 7B) and survivor fish (Fig 7C), respectively, when compared to that of control fish. However, in serum (Fig 7D–F), parasite-specific IgT binding was detected only in 1/10 dilution of survivor fish (Fig 7F). In contrast, parasite-specific IgM binding in up to 1/1000 and 1/4000 of the diluted serum of infected (Fig 7E) and survivor fish (Fig 7F) increased by ~ 2.9-fold and ~ 4.3-fold, respectively. Finally, in the nasal mucosa and serum of both the infected and survivor fish, Ich-specific IgD showed no change when compared to that of control fish (Fig 7A–F).

**Fig 7.**
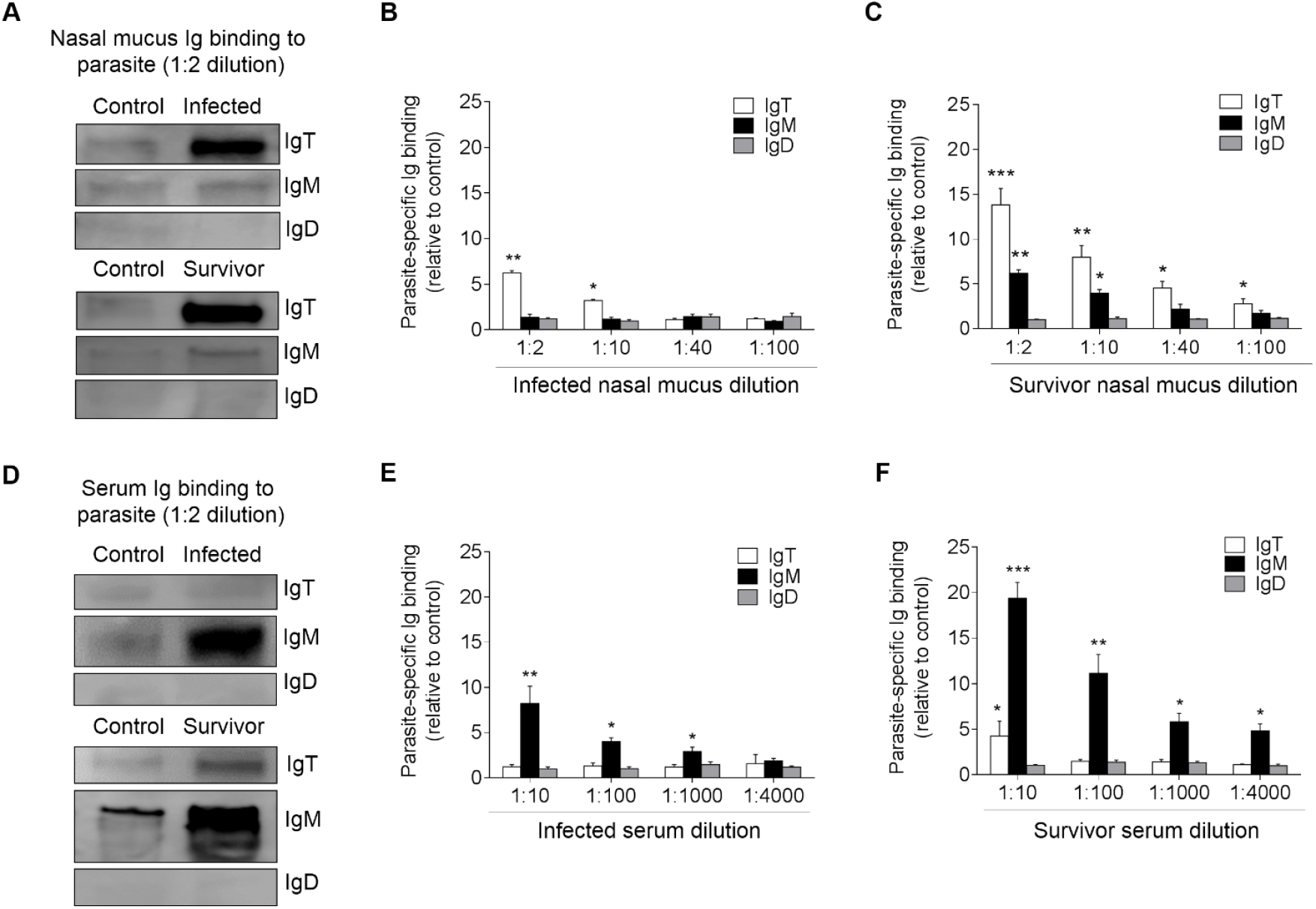
Immunoglobulin responses in the nasal mucus and serum from infected and survived trout. (A) Western blot analysis of IgT-, IgM- and IgD-specific binding to Ich in nasal mucus (dilution 1:2) from infected and survivor fish. (B and C) IgT-, IgM- and IgD-specific binding to Ich in dilutions of nasal mucus from infected (B) and survivor (C) fish, evaluated by densitometric analysis of immunoblots and presented as relative values to those of control fish (n = 8 per group). (D) Western blot analysis of IgT-, IgM- and IgD-specific binding to Ich in serum (dilution 1:10) from infected and survivor fish. (E and F) IgT-, IgM- and IgD-specific binding to Ich in dilutions of serum from infected (E) and survivor (F) fish, evaluated by densitometric analysis of immunoblots and presented as relative values to those of control fish (n = 8 per group). \**P* < 0.05, \*\**P* < 0.01 and \*\*\**P* < 0.001 (unpaired Student’s *t*-test). Data are representative of at least three independent experiments (mean and s.e.m.).

### Local proliferation of B-cells and Ig responses in trout olfactory organ after Ich parasite infection

To further evaluate whether an increase of IgT^+^ B-cells in the olfactory organ of survivor fish was derived from the process of local IgT^+^ B-cell proliferation or from influx of B cells from systemic lymphoid organs, we performed *in vivo* proliferation studies of IgT^+^ B-cells and IgM^+^ B-cells stained with EdU, which can incorporate into DNA during cell division [26]. Immunofluorescence microscopy analysis showed a significant increase in the percentage of proliferating cells in the olfactory organ of survivor fish (~ 0.048 ± 0.0006 %) when compared with that of control animals (~ 0.019 ± 0.0003 %) (Fig 8A and B). Interestingly, we detected a significant increase in the proliferation of EdU^+^ IgT^+^ B-cells in survivor fish (~ 5.21 ± 0.23 %) when compared with that of the control fish (~ 0.58 ± 0.05 %) (Fig 8A and C). However, no difference was found in the percentage of EdU^+^ IgM^+^ B-cells of control fish and survivor fish (Fig 8A–C). Using flow cytometry, similar results were obtained (S4A Fig), with the percentage of EdU^+^ IgT^+^ B-cells increased significantly in the olfactory organ of survivor fish (~ 5.67 ± 0.10% in all IgT^+^ B-cells) when compared with that of control fish (~ 3.46 ± 0.26% in all IgT^+^ B-cells), while no difference in the percentage of EdU^+^ IgM^+^ B-cells was detected between control and survivor fish (S4A Fig). In the head kidney, the percentage of EdU^+^ IgM^+^ B-cells of the olfactory organ was detected in survivor fish, and it presented a large increase when compared with that of control fish. In contrast, these two groups showed no difference in proliferating IgT^+^ B-cells (S4B Fig). The local proliferation of IgT^+^ B-cells in the olfactory organ and the detection of parasite-specific IgT in nasal mucus (Fig 7) suggest that specific IgT in the trout olfactory organ is locally generated rather than produced and transported from systemic lymphoid organs.

**Fig 8.**
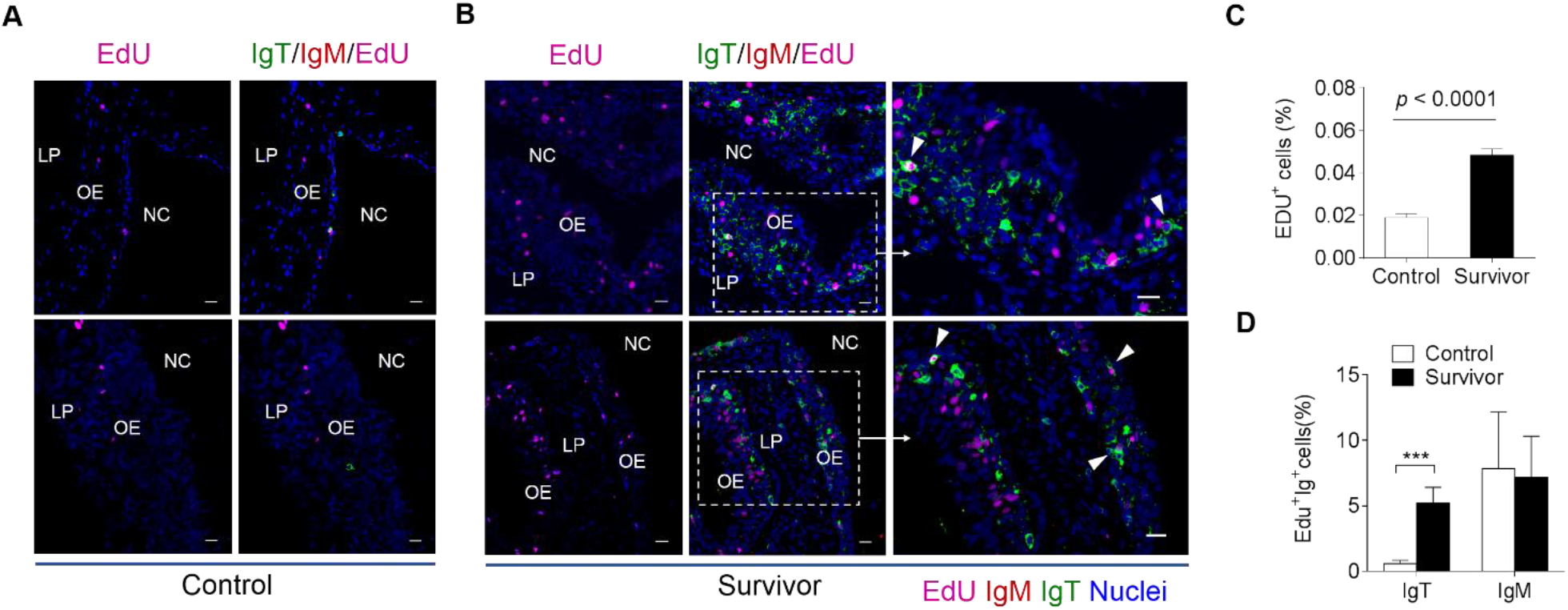
Proliferative responses of IgT^+^ and IgM^+^ B cells in the olfactory organ of survived trout. Immunofluorescence analysis of EdU incorporation by IgT^+^ or IgM^+^ B cells in olfactory organ of control (A) and survivor fish (B). Nasal paraffin sections were stained for EdU (magenta), trout IgT (green), trout IgM (red) and nuclei (blue) detection (*n* = 8 fish per group). NC, nasal cavity; OE, olfactory epithelium; LP, lamina propria. Scale bars, 20 μm. (C) Percentage of EdU^+^ cells from total nasal cell in control or survivor fish counted from Fig 7A and B (*n* = 8). (D) Percentage of EdU^+^ cells from the total IgT^+^ or IgM^+^ B cells populations in olfactory organ of control and survivor fish counted from A and B. Data in A and B are representative of at least three independent experiments (mean and s.e.m.). Statistical analysis was performed by unpaired Student’s *t*-test. \**P* < 0.05, *\*\**P** < 0.01 and \*\*\**P* < 0.001.

To further address this hypothesis, we measured parasite-specific Igs titers from medium of cultured olfactory organ, head kidney, and spleen explants from control and survivor fish (Fig 9). We detected parasite-specific IgT binding in 1/40 diluted medium (~ 3.6-fold) of cultured olfactory organ explants of survivor fish, whereas low parasite-specific IgM titers were detected only at the 1/10 dilution in the same medium (Fig 9A and D). In contrast, dominant parasite-specific IgM binding (up to 1/40 dilutions) was observed in the medium of head kidney and spleen explants, and low parasite-specific IgT responses were detected in the same medium (Fig 9B–F). Interestingly, negligible parasite-specific IgD titers were detected in the medium of cultured olfactory organ, head kidney, and spleen explants from control and survivor fish (Fig 9A–F). Combined, these results demonstrate that specific Ig responses against parasites are compartmentalized in rainbow trout with IgT present in the olfactory organ and IgM present in systemic lymphoid tissues and serum.

**Fig 9.**
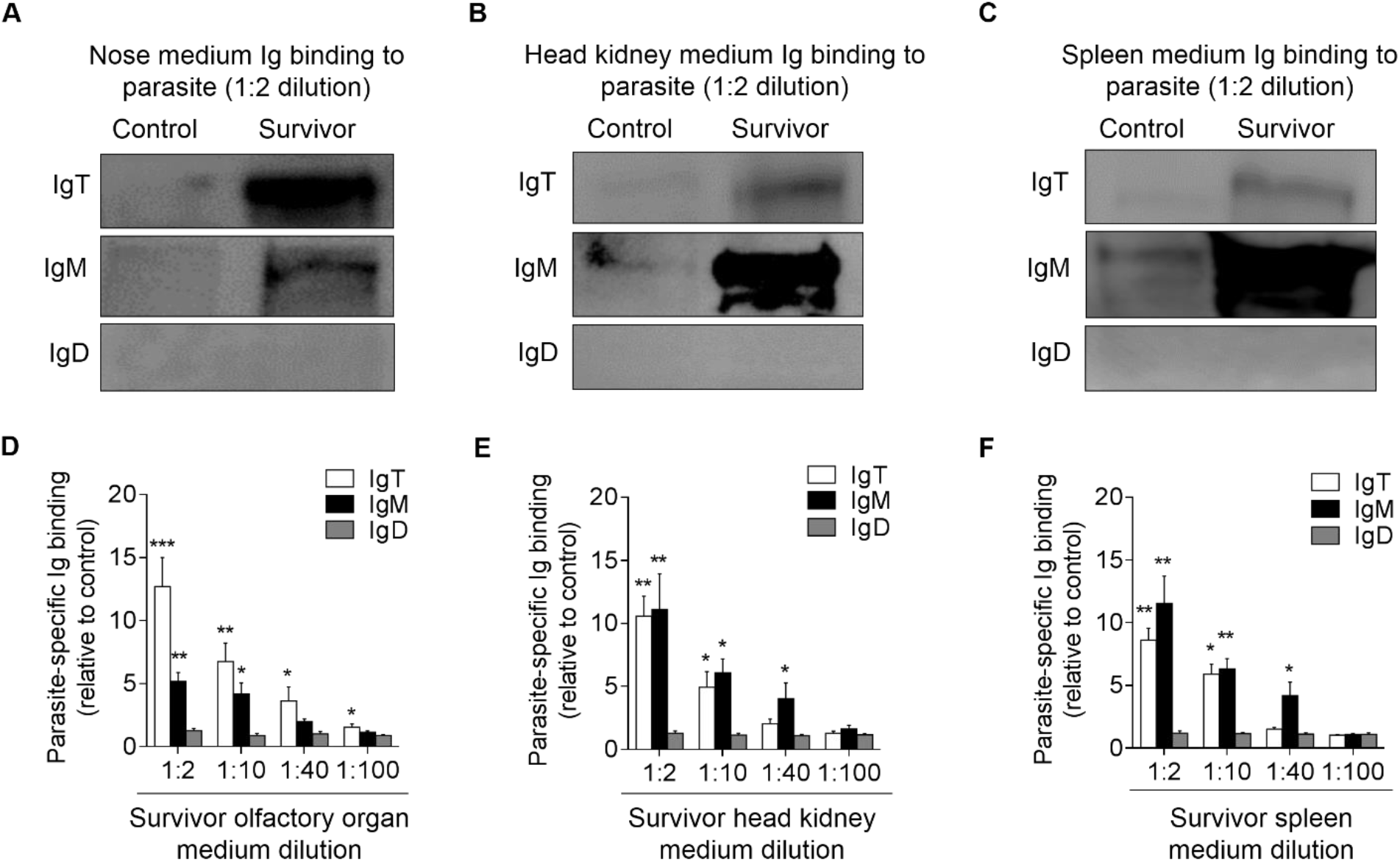
Local IgT-, IgM- and IgD-specific responses in olfactory organ explants of survivor fish. The olfactory organ, head kidney and spleen explants (~ 20 mg each) from control and survivor fish were cultured for 7 days. Immunoblot analysis of IgT-, IgM- and IgD-specific binding to Ich in the culture medium of olfactory organ (A), head kidney (B) and spleen (C) (dilution 1:2) from control and survivor fish. (D-F) IgT-, IgM- and IgD-specific binding to Ich in dilutions of culture medium from olfactory organ (D), head kidney (E) and spleen (F) from control and survivor fish, measured by densitometric analysis of immunoblots and presented as relative values to those of control fish (*n* = 6-8 per group). \**P* < 0.05, \*\**P* < 0.01 and \*\*\**P* < 0.001 (unpaired Student’s *t*-test). Data are representative of at least three independent experiments (mean and s.e.m.).

## Discussion

Protozoans are the most common parasites of freshwater and marine fish [29–31]. Ich is one of the most problematic parasites in freshwater ecosystems infecting many different fish species [32, 33]. Ich has been traditionally associated with skin and gill lesions in rainbow trout [15, 17], however, the teleost olfactory organ is constantly exposed to the aquatic environment and therefore may represent a route of entry for any pathogen. Here we report for the first time that Ich can infect the olfactory organ of rainbow trout when the fish are exposed to the parasite by bath, the natural route of exposure. Importantly, we found that parasite loads in the olfactory organ were the highest along with the gills, suggesting that the olfactory route of infection may be one of the main targets of this parasite. Moreover, Ich transcript levels were detectable in the olfactory organ up to 75 days post exposure, indicating that Ich establishes long-term infections in this tissue. Given that the impacts of Ich invasion via the nose have until now been overlooked, further investigations are required to determine the impacts of Ich nasal infections in the fish host health.

The olfactory organ of teleosts, similar to that of mammals, is coated by mucus containing Igs. In this study, we characterized in detail all three Ig classes in the nasal mucus of rainbow trout, including sIgD, secreted IgM (sIgM), and secreted IgT (sIgT). Trout nasal IgT existed for the most part as a polymer, similar to the characterized IgA in the nasal mucosa from humans [34, 35]. On the contrary, nasal IgD was in monomeric form, as previously reported in the gill [15] and serum [36]. Interestingly, in agreement with the descriptions for gut [16], skin [17], and gill [15] sIgT, all subunits of polymeric nasal IgT in rainbow trout were associated by noncovalent interactions. In addition, we detected the concentrations of all three Igs in nasal mucosa and serum and found that although the concentration of IgT was lower than that of IgM in both nasal mucosa and serum, the ratio of IgT/IgM in nasal mucosa was higher than that in serum, in agreement with a previous report by Tacchi et al [9]. Combined, these findings underscore that mucus secretions in teleosts consist of mixtures of all three Ig isotypes and that Ig protein concentrations of each isotype differ among the four teleost MALT [15–17] as they do in mammals [37–39].

In mammals, NALT has been considered a mucosal inductive site for IgA [40–42]. Yet, it is not clear whether in fish, which lack organized lymphoid structures (adenoids and tonsils) in the teleost olfactory organ [9], NALT acts as an inductive and/or effector mucosal lymphoid tissue. In our Ich infection model, we found large increases in the concentration of IgT but not IgM or IgD at the protein level in the nasal mucosa of infected and surviving fish exposed to Ich, which correlated with the large accumulation of IgT^+^ but not IgM^+^ B-cells appearing in the olfactory epidermis of the same fish. In support, we showed a striking abundance of IgT coating on the Ich parasite surface in the olfactory organ of rainbow trout. However, much lower or negligible levels of IgM or IgD coating were detected on the same parasites. These results suggested that a strong IgT but not IgM response to Ich takes place in the local olfactory environment. Interestingly, similar results were also discovered in our previous studies in the gut, skin, and gills [15–17]. In mammals, a dramatic increase of IgA secretion and significant accumulations of IgA-antibody forming cells (IgA-AFC) were induced in the nasal mucosa following intranasal infection with a small volume of influenza virus [18] and *N. fowleri* parasite [43–45], respectively. Based on our findings, it is clear that teleost NALT is a mucosal inductive site. Whether NALT-induced IgT^+^ B-cell and plasma cell responses seed effector sites such as the gut lamina propria remains to be characterized in this or other models. Finally, our results strengthen the notion that despite anatomical differences and the absence of organized NALT structures in teleosts, IgT and IgA carry out play vital roles in nasal adaptive immune responses.

Immunoglobulins are of particular relevance in the context of Ich infections since previous studies have demonstrated that antibody (IgM) mediated responses against Ich i-antigen trigger the exit of the parasite from the fish host skin conferring host disease resistance [46, 47]. Similarly, in our model, Ich was being expulsed at day 28 and minority stay in nasal cavity, but interestingly, expelled Ich was mainly coated by IgT. In addition, we recorded the greatest upregulation in the expression of the IgT heavy chain gene in the trout olfactory organ 28 days after Ich exposure, the same time point when Ich levels dropped dramatically, suggesting that IgT might play a crucial role in the nasal immune response to Ich infection and may contribute to parasite clearance or exit. At this point, IgM expression levels had also increased in the olfactory organ and some detectable titers of parasite-specific IgM were found in trout nasal mucosa. Nasal IgM titers might be the result of Ich-instigated microlesions in the olfactory system and consequent leakage of parasite-specific IgM or plasma from the blood. Thus, specific IgT responses appear to be the most critical antibody response against Ich in the nasal environment and further studies should address how IgT contributes to parasite clearance from trout mucosal surfaces.

Interestingly, similar to the previous results in the gill, our results indicated that negligible parasite-specific IgD responses were induced in both the nasal mucosa and serum after Ich challenges. However, because of the detectable concentrations of IgD in both the nasal mucosa and serum, we cannot exclude the possibility that relevant IgD may be induced in the nasal mucosa or systemic compartment when using different pathogens or stimulation routes. Thus, future studies are needed to investigate the role of nasal and systemic IgD in the parasite-specific immune responses of teleost fish against different pathogens.

The accumulations of IgT^+^ B-cells observed in the olfactory epidermis correlated with high parasite-specific IgT titers in the same fish led us to hypothesize the local proliferation and production of the parasite-specific IgT^+^ B-cell response. IgT^+^ B-cell proliferation responses were detected in the olfactory organ but not in the systemic immune organs (head kidney and spleen) of the same fish, which strongly suggests that the accumulation of IgT^+^ B-cells in the olfactory organ is due to local proliferation rather than migration from other organs. We also show that olfactory tissue explants produce specific anti-Ich IgT antibodies, demonstrating the presence of specific plasma cells in the local nasal mucosa. Interestingly, these results parallel our previous finding in the trout gill, and the proliferation rates we detected here was similar to the ones we have described in gills [15] but higher than those in olfactory organ in response to IHNV [21], which might be due to the different duration after infection/immunization with different pathogen, respectively. It is worth noting that similar results were found in the NALT of mammals. For instance, previous studies have shown that intranasal immunization with *Naegleria fowleri* could induce the secretion of IgA and IgG in nasal mucosa but pathogen-specific IgA mainly mediates local nasal immunity in mammals [48, 49]. By *in vitro* culture of NALT cells following virus infection, parts of the virus-specific antibody-forming cells (AFCs) were observed to originate from B-cell precursors in NALT [18]. Moreover, in the nasal mucosa from 53 humans with chronic inflammation, most IgA seemed to be produced locally by IgA-producing plasma cells [50]. Hence, our results indicate that the local proliferation of mucosal B-cells and production of secretory Ig responses in the nasal mucosa happens not only in tetrapod species but also in early vertebrates such as teleost fish.

The fish olfactory mucosa is a complex neuroepithelium in which lymphoid and myeloid cells are found in a scattered manner [9] among basal cells, sustentacular cells, olfactory sensory neurons, goblet cells and epithelial cells (Figure 10). In agreement with histological changes, strong immune responses including the upregulation of cytokine expression and complement genes were detected in the olfactory organ, especially at the early stages of the infection, preceding the onset of Ig responses. Interestingly, in mammals [43], intranasal administration with *N. fowleri* lysates plus cholera toxin (CT) results in increased expression of genes for IL-10, IL-6, IFN-γ, TNF-α, and IL-1β. This immune expression signatures largely resemble those found in the present study. Thus, despite the lack of organized lymphoid structures in the olfactory organs of teleosts [3, 9], teleost fish mount strong cytokine responses upon pathogen invasion or immunization with antigenic analogues.

**Fig 10.**
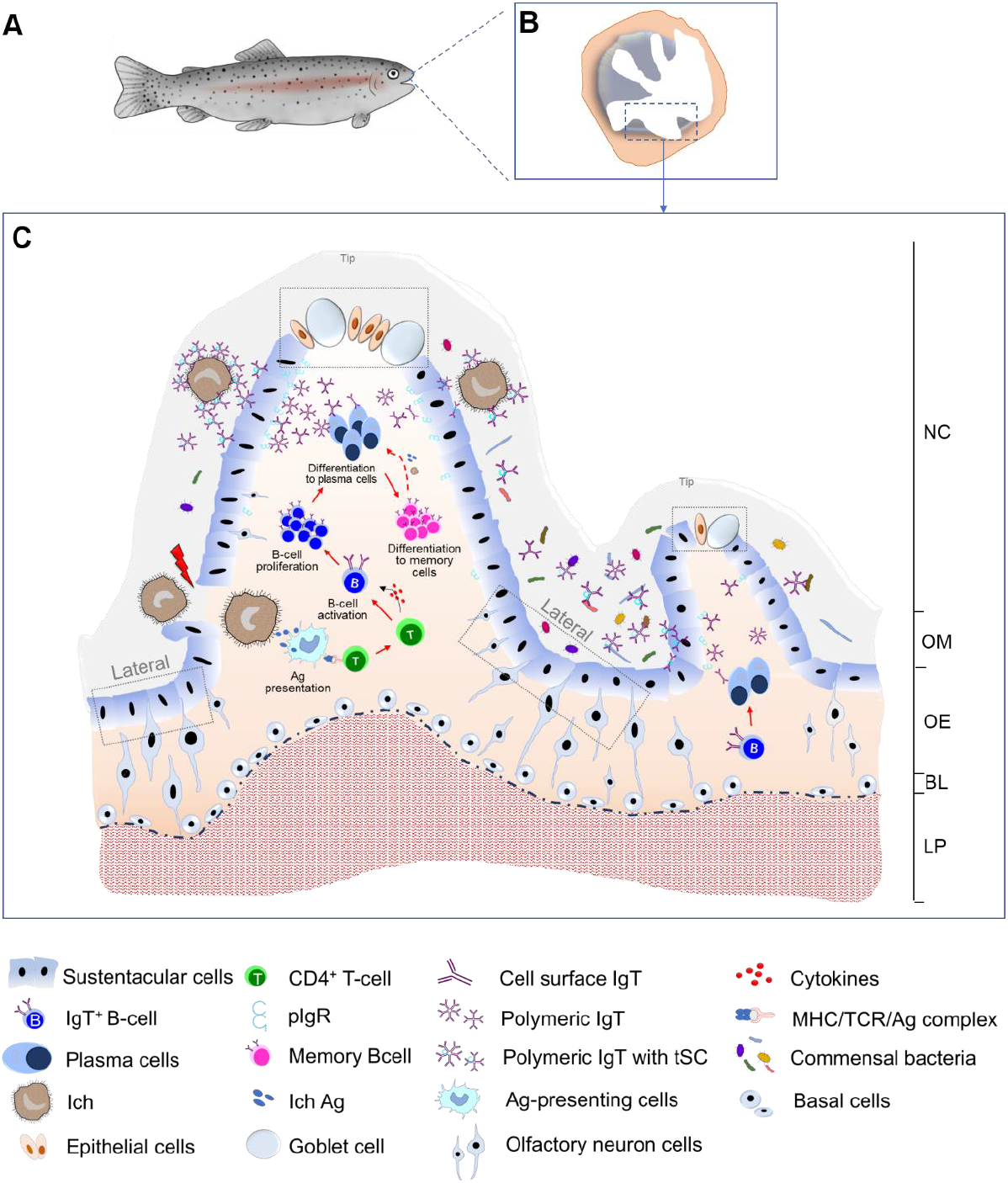
Proposed model of local IgT and IgT^+^ B cell induction in the olfactory organ. Model images represent a fish (A) and enlarged sections of the areas outlined in a showing the olfactory organ (B). (C) Induction of local IgT responses in the trout olfactory organ based on our findings. When Ich parasite invaded the nasal mucosa, Ich antigen (Ag) are taken up by antigen-presenting cells (APC) and presented to naïve CD4-T cells. Ag-specific CD4-T cells then produced cytokines to activate B cells. Activated B cells start proliferating in olfactory organ and may differentiation to plasma cells to locally produce Ich-specific IgT, which will be transported by pIgR into nasal mucus where can specific binding to the Ich parasite. Alternatively, some IgT^+^ plasma cells may differentiate into memory IgT^+^ B cells. When Ich parasite infection happened again in olfactory organ, memory IgT^+^ B cells directly proliferate and differentiate into plasma cells and produce larger amounts of specific-IgT to binding Ich. The trout olfactory organ showing the mucosal tip area with goblet cells and the lateral neuroepithelium. NC, nasal cavity; OM, olfactory mucus; OE, olfactory epithelium; LP, lamina propria.

In conclusion, our results provide the first evidence that parasite infection, antigen presentation, local B-cell activation and proliferation, as well as parasite-specific IgT production occur in the olfactory organ of teleost fish (Fig 10). Thus, although parasites such as Ich can infect the olfactory organ of fish, local IgT^+^ B-cells and parasite-specific IgT appear to be a major mechanism by which the host acquires resistance to this parasite. Our findings not only expand our view of nasal immune systems from an evolutionary perspective but also suggest that nasal vaccination may be an effective way to prevent aquatic parasitic diseases.

## Materials and methods

### Ethics statement

All experimental protocols involving fish were performed in accordance with the recommendations in the Guide for the care and use of Laboratory Animals of the Ministry of Science and Technology of China and were approved by the Scientific Committee of Huazhong Agricultural University (permit number HZAUFI-2016-007).

### Fish

Rainbow trout (20-30 g) were obtained from fish farm in Shiyan (hubei, China), and maintained them in aquarium tanks using a water recirculation system involving thermostatic temperature control and extensive biofiltration. Fish were acclimatized for at least 2 wk at 15 °C and fed daily with commercial trout pellets at a rate of 0.5-1 % body weight day^−1^ and feeding was terminated 48 h prior to sacrifice.

### Ich parasite isolation and infection

The method used for Ich parasite isolation and infection were described previously by Xu et al [17] with slight modification. Briefly, heavily infected rainbow trout were anaesthetized with overdose of MS-222 and placed in a beaker with water to allow trophonts and tomonts exit the fish. The trophonts and tomonts were left in the water at 15 °C for 24 h to let tomocyst formation and subsequent theront release. For parasite infection, two types of challenges with Ich were performed. The first group, fish were exposed to a single dose of ~ 5,000 theronts per fish added into the aquarium, and tissue samples and fluids (serum and nasal mucus) were taken after 28 days (infected fish). The second group, fish were monthly exposed during 75 days period with ~ 5,000 theronts per fish (survival fish). Fish samples were taken two weeks after the last challenge. Experiments were performed at least three independent times. Control fish (mock infected) were maintained in a similar tank but without parasites. During the whole experiment periods, the fish were raised in a flow through aquaria at 15 °C and fed daily with commercial trout pellets at a rate of 0.5-1 % body weight day^−1^.

### Collection of serum, olfactory tissue and nasal mucus

For sampling, trout were anaesthetized with MS-222 and serum was collected and stored as described [16]. To obtain fish nasal mucus, we modified the method described previously [9, 15]. Briefly, trout olfactory tissue was excised rinsed with PBS three times to remove the remaining blood. Thereafter, olfactory tissue was incubated for 12 h at 4 °C, with slightly shaking in protease inhibitor buffer (1 × PBS, containing 1 × protease inhibitor cocktail (Roche), 1 mM phenylmethylsulfonyl fluoride (Sigma); pH 7.2) at a ratio of 100 mg of olfactory tissue per ml of buffer. The suspension (nasal mucus) was transferred to an Eppendorf tube, and then the supernatant was vigorously vortexed and centrifuged at 400 *g* for 10 min at 4 °C to remove trout cells. Furthermore, the olfactory organ was taken and fixed into 4 % neutral buffered formalin for hematoxylin and eosin (H & E) staining and immunostaining.

### Isolation of trout HK and NALT leukocytes

The leucocytes from head kidney were obtained using a modified methodology as described previously [15, 24]. To obtain trout nasopharynx-associated lymphoid tissue (NALT) leukocytes, we modified the existing protocol as explained by Tacchi et al [9]. Briefly, rainbow trout were anaesthetized with MS-222 and blood was collected from the caudal vein. The olfactory organ was taken and washed with cold PBS to avoid blood contamination. Leucocytes from trout olfactory organ were isolated by mechanical agitation of both olfactory rosettes in DMEM medium (supplemented with 5 % FBS, 100 U ml ^−1^ penicillin and 100 μg ml ^−1^ streptomycin) at 4 °C for 30 min with continuous shaking. Leukocytes were collected, and the aforementioned procedure was repeated four times. Thereafter, the olfactory organ pieces were treated with PBS (containing 0.37 mg ml ^−1^ EDTA and 0.14 mg ml ^−1^ dithiothreitol DTT) for 30 min followed by enzymatic digestion with collagenase (Invitrogen, 0.15 mg ml ^−1^ in PBS) for 1 h at 20 °C with continuous shaking. All cell fractions obtained from the olfactory organ after mechanical and enzymatic treatments were washed three times in fresh modified DMEM and layered over a 51/34 % discontinuous Percoll gradient. After 30 min of centrifugation at 400 *g*, leucocytes lying at the interface of the gradient were collected and washed with modified DMEM medium.

### SDS-PAGE and western blot

Nasal mucus and serum samples were resolved on 4-15 % SDS-PAGE Ready Gel (Bio-Rad) under non-reducing or reducing conditions as described previously [15–17]. For western blot analysis, the gels were transferred onto PVDF membranes (Bio-Rad). Thereafter, the membranes were blocked with 8 % skim milk and incubated with anti-trout IgT (rabbit pAb), anti-trout IgM (mouse monoclonal antibody (mAb)) or biotinylated anti-trout IgD (mouse mAb) antibodies followed by incubation with peroxidase-conjugated anti-rabbit, anti-mouse IgG (Invitrogen) or streptavidin (Invitrogen). Immunoreactivity was detected with an enhanced chemiluminescent reagent (Advansta) and scanned by GE Amersham Imager 600 Imaging System (GE Healthcare). The captured gel images were analysed by using ImageQuant TL software (GE Healthcare). Thereafter, the concentration of IgM, IgD and IgT were determined by plotting the obtained signal strength values on a standard curve generated for each blot using known amounts of purified trout IgM, IgD or IgT.

### Gel filtration

To analysis the monomeric or polymeric state of Igs in trout nasal mucus, gel filtration analyses were performed using as described previously for gut [16] and gill mucus [15]. In short, fractions containing the IgM, IgT or IgD were separated by gel filtration using a Superdex-200 FPLC column (GE Healthcare). The column was previously equilibrated with cold PBS (pH 7.2), and protein fractions were eluted at 0.5 ml min ^−1^ with PBS using a fast protein LC instrument with purifier systems (GE Healthcare). Identification of IgM, IgD and IgT in the eluted fractions was performed by western blot analysis using anti-IgM, anti-IgD and anti-IgT antibodies, respectively. A standard curve was generated by plotting the elution volume of the standard proteins in a Gel Filtration Standard (Bio-Rad) against their known molecular weight, which was then used to determine the molecular weight of the eluted IgT, IgM and IgD by their elution volume.

### Flow cytometry

For flow cytometry studies of B cells in the head kidney and NALT, leukocyte suspensions were double-stained with monoclonal mouse anti-trout IgT and anti-trout IgM (1 μg ml ^−1^ each) at 4°C for 45 min. After washing three times, PE-goat anti-mouse IgG1 and APC-goat anti-mouse IgG2b (1 μg ml ^−1^ each, BD Biosciences) were added and incubated at 4 °C for 45 min to detect IgM^+^ and IgT^+^ B cells, respectively. After washing three times, analysis of stained leucocytes was performed with a CytoFLEX flow cytometer (Beckman coulter) and analysed by FlowJo software (Tree Star).

### Histology, light microscopy and immunofluorescence microscopy studies

The olfactory organ of rainbow trout was dissected and fixed in 4 % neutral buffered formalin overnight at 4 °C and then transferred to 70 % ethanol. Samples were then embedded in paraffin and 5 μm thick sections stained with haematoxylin / eosin (H & E). Images were acquired in a microscope (Olympus) using the Axiovision software. For the detection of Ich parasite at the same time of IgT^+^ and IgM^+^ B cells, sections were double-stained with rabbit anti-trout IgT (pAb; 0.49 μg ml^−1^) and mouse anti-trout IgM (IgG1 isotype; 1 μg ml^−1^) overnight at 4°C. After washing three times, secondary antibodies Alexa Fluor 488-conjugated AffiniPure Goat anti-rabbit IgG or Cy3-conjugated AffiniPure Goat anti-mouse IgG (Jackson ImmunoResearch Laboratories Inc.) at 2.5 μg ml^−1^ each were added and incubated at temperature for 40 min to detect IgT^+^ and IgM^+^ B cells, respectively. After washing three times, mouse anti-Ich polyclonal antibody (1 μg ml^−1^) were added and incubated at 4 °C for 6 h, after washing three times, secondary antibody Alexa Fluor 647-goat anti-mouse (Jackson ImmunoResearch Laboratories Inc.) with 5 μg ml^−1^ were added and incubated at temperature for 40 min to detect Ich parasite. For detection of trout nose pIgR, we used the same methodology described to stain gill pIgR by using our rabbit anti-pIgR [16]. As controls, the rabbit IgG pre-bleed and the mouse-IgG1 isotype antibodies were labelled with the same antibody labelling kits and used at the same concentrations. Before mounting, all samples were stained with DAPI (4’, 6-diamidino-2-phenylindole; 1 μg ml^−1^: Invitrogen) for the sections. Images were acquired and analysed using Olympus BX53 fluorescence microscope (Olympus) and the iVision-Mac scientific imaging processing software (Olympus).

### Proliferation of B cells in the olfactory organ of trout

For proliferation of B cells studies, we modified the methodology as previously reported by us [15]. Briefly, control and survivor fish (~ 30g) were anaesthetized with MS-222 and intravenous injected with 200 μg EdU (Invitrogen). After 24 h, leucocytes from head kidney or olfactory tissue were obtained as described above, and cells were incubated with 10 μM of EdU (Invitrogen) for 2 hours. Thereafter, leucocytes were incubated with mAb mouse anti-trout IgM and anti-trout IgT (1 μg ml^−1^ each) at 4 °C for 45 min. After washing three times, Alexa Fluor 488-goat anti-mouse IgG (Invitrogen) was used as secondary antibody to detect IgM^+^ or IgT^+^ B cells. After incubation at 4 °C for 45 min, cells were washed three times with DMEM medium and fixed with 4 % neutral buffered formalin at room temperature for 15 min. EdU^+^ cell detection was performed according to the manufacturer’s instructions (Click-iT EdU Alexa Fluor 647 Flow Cytometry Assay Kit, Invitrogen). Cells were thereafter analysed in a CytoFLEX flow cytometer (Beckman coulter) and FlowJo software (Tree Star). For immunofluorescence analysis, as described above, we used the paraffin sections of olfactory organ from control and survival fish previously injected with EdU and incubated at 4 °C for 45 min with rabbit anti-trout IgT (pAb; 1 μg ml^−1^) and mouse anti-trout IgM (IgG1 isotype; 1 μg ml^−1^). After washing with PBS, paraffin sections were incubated for 2 h at room temperature with Alexa Fluor 488-conjugated AffiniPure Goat anti-rabbit IgG or Cy3-conjugated AffiniPure Goat anti-mouse IgG (Jackson ImmunoResearch Laboratories Inc.) at 2.5 μg ml^−1^ each. Stained cells were fixed with 4 % neutral buffered formalin and EdU^+^ cell detection was performed according to the manufacturer’s instructions (Click-iT EdU Alexa Fluor 647 Imaging Kit, Invitrogen). Cell nuclei were stained with DAPI (1 μg ml^−1^) before mounting with fluorescent microscopy mounting solution. Images were acquired and analysed using an Olympus BX53 fluorescence microscope (Olympus) and the iVision-Mac scientific imaging processing software (Olympus).

### Tissue explants culture

To assess whether the parasite-specific IgT responses were locally generated in the olfactory organ, we analysed parasite-specific immunoglobulin titers from medium derived of cultured olfactory organ, head kidney and spleen explants obtained from control and survivor fish as previously described by us [15]. In short, control and survivor fish were anaesthetized with an overdose of MS-222, and blood was removed through the caudal vein to minimize the blood content in the collected organs. Thereafter, olfactory organ, head kidney and spleen were collected. Approximately 20 mg of each tissue was submerged in 70 % ethanol for 1 min to eliminate possible bacteria on their surface and then washed twice with PBS. Thereafter, tissues were placed in a 24-well plate and cultured with 200 ml DMEM medium (Invitrogen), supplemented with 10 % FBS, 100 U ml ^−1^ penicillin, 100 μg ml ^−1^ streptomycin, 200 μg ml ^−1^ amphotericin B and 250 μg ml ^−1^ gentamycin sulfate, with 5 % CO_2_ at 17 °C. After 7 days culture, supernatants were harvested, centrifuged and stored at 4 °C prior to use the same day.

### Binding of trout immunoglobulins to Ich

The capacity of IgT, IgM and IgD from serum, nasal mucus or tissue (olfactory organ, head kidney and spleen) explant supernatants to bind to Ich was measured by using a pull-down assay as described previously [15, 17]. Briefly, approximately 100 tomonts were pre-incubated with a solution of 0.5 % BSA in PBS (pH 7.2) at 4 °C for 2 h. Thereafter, tomonts were incubated with diluted nasal mucus or serum or tissue (olfactory organ, head kidney and spleen) explant supernatants from infected, survivor or control fish at 4 °C for 4 h with continuous shaking in a 300 ml volume. After incubation, the tomonts were washed three times with PBS and bound proteins were eluted with 2 × Laemmli Sample Buffer (Bio-Rad) and boiled for 5 min at 95 °C. The eluted material was resolved on 4-15 % SDS-PAGE Ready Gel under non-reducing conditions, and the presence of IgT, IgM or IgD was detected by western blotting using anti-trout IgT, IgM or IgD antibodies as described above.

### RNA isolation and quantitative real-time PCR (qPCR) analysis

Total RNA was extracted by homogenization in 1 ml TRIZol (Invitrogen) using steel beads and shaking (60 HZ for 1 min) following the manufacturer’s instructions. The quantification of the extracted RNA was carried out using a spectrophotometry (NanoPhotometer NP 80 Touch) and the integrity of the RNA was determined by agarose gel electrophoresis. To normalize gene expression levels for each sample, equivalent amounts of the total RNA (1000 ng) were used for cDNA synthesis with the SuperScript first-strand synthesis system for qPCR (Abm) in a 20 μl reaction volume. The synthesized cDNA was diluted 4 times and then used as a template for qPCR analysis. The qPCRs were performed on a 7500 Real-time PCR system (Applied Biosystems) using the EvaGreen 2 × qPCR Master mix (Abm). All samples were performed following conditions: 95 °C for 30 s, followed by 40 cycles at 95 °C for 1 s and at 58 °C for 10 s. A dissociation protocol was carried out after thermos cycling to confirm a band of the correct size was amplified. Ct values determined for each sample were normalized against the values for housekeeping gene (EF1α). To gain some insights on the kinetics of the immune responses that takes place after Ich infection, twenty-six immune relevant genes, such as cytokine, complement and Igs genes were detected in the olfactory organ. The relative expression level of the genes was determined using the Pfaffl method [25]. The primers used for qRT-PCR are listed in Sup1 Table.

### Co-immunoprecipitation studies

We followed the same strategy to detect the association of pIgR to IgT in gut, skin and gill mucus as we previously described [15–17]. To detect whether polymeric trout IgT present in the nasal mucus were associated to a secretory component-like molecule derived from trout secretory component–like molecule (tSC), we performed co-immunoprecipitating analysis using anti-trout IgT (pAb) antibodies with the goal to potentially co-immunoprecipitate the tSC. To this end, 10 μg of anti-IgT were incubated with 300 μl of trout nasal mucus. As control for these studies, the same amount of rabbit IgG (purified from the pre-bleed serum of the rabbit) were used as negative controls for anti-IgT. After overnight incubation at 4°C, Dynabeads Protein G (10001D; 50μl; Invitrogen) prepared previously was added into each reaction mixture and incubated for 1 h at 4 °C following the manufacturer’s instructions. Thereafter, the beads were washed five times with PBS, and subsequently bound proteins were eluted in 2 × Laemmli Sample Buffer (Bio-Rad). The eluted material was resolved by SDS-PAGE on 4–15% Tris-HCl Gradient ReadyGels (Bio-Rad) under reducing (for tSC detection) or non-reducing (for IgT detection) conditions. Western blot was performed with anti-pIgR and anti-IgT antibodies as described above.

### Statistical analysis

An unpaired Student’s t-test and one-way analysis of variance with Bonferroni correction (Prism version 6.01; GraphPad) were used for analysis of differences between groups. Data are expressed as mean ± s.e.m. *P* values less than 0.05 were considered statistically significant.

## Acknowledgments

We thank Dr. J. Oriol Sunyer (University of Pennsylvania) for his generous gift of anti-trout IgM, anti-trout IgD, anti-trout IgT mAbs, anti-trout IgT and anti-trout pIgR pAbs.

## Supporting information

**S1 Fig.**
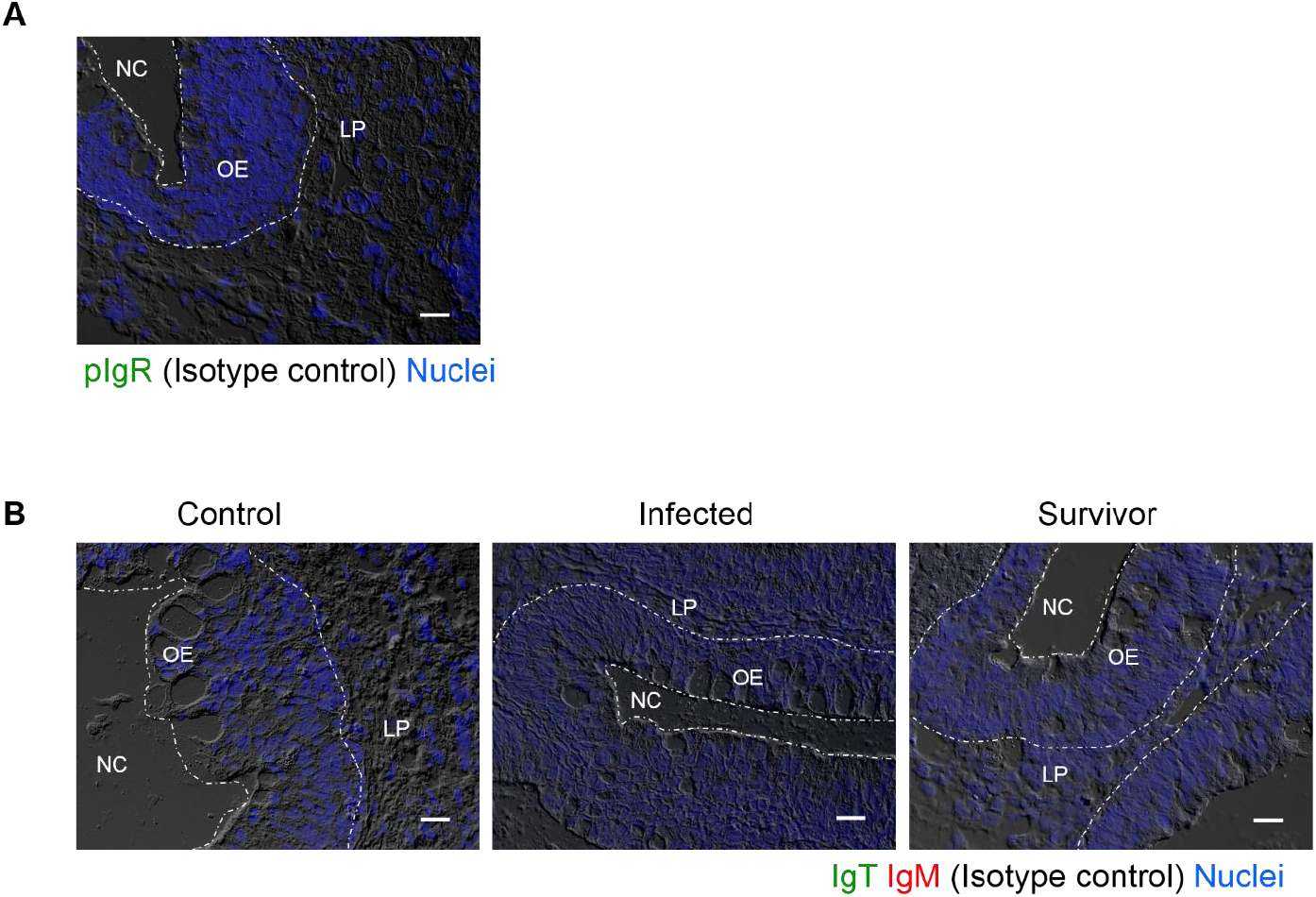
Isotype control staining for anti-IgT, anti-IgM and anti-pIgR antibodies in trout olfactory organ paraffin sections. Differential interference contrast images of olfactory organ paraffin sections from 28 days Ich-infected fish (A middle and B), survivor fish (A right), and control fish (A left), with merged staining of isotype control antibodies for anti-trout IgT (green) or anti-trout IgM mAbs (red) (A); or for anti-trout pIgR pAb (green, B). Nuclei were stained with DAPI (blue, A and B). NC, nasal cavity; OE, olfactory epithelium; LP, lamina propria. Scale bar, 20 μm. Data are representative of at least three different independent experiments.

**S2 Fig.**
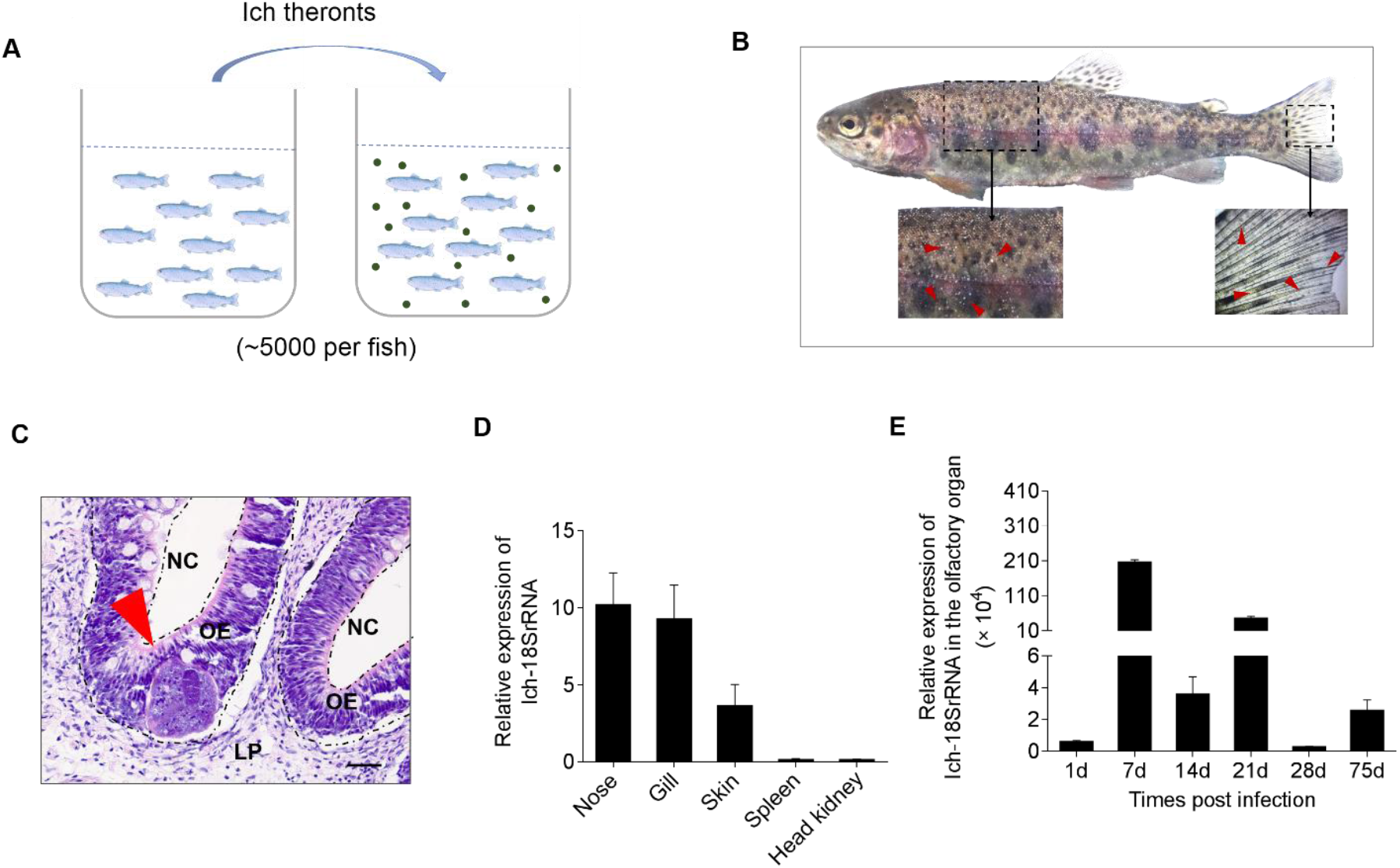
The detection of Ich parasite in olfactory organ of trout infected with Ich. (A) Infection method with Ich parasite by bath has been used in this study. (B) The phenotype of rainbow trout was observed at 7 days post infection with Ich (*n* = 12). The red arrows represent the obvious while dot in skin (lower, left) and fin (lower, right). (C) Histological studies of olfactory organ from 7 days Ich-infected trout by staining with haematoxylin / eosin (H & E). Results are representative of one experiment *n* = 6. Scale bar: 50 μm. (D) The relative expression of Ich-18SrRNA gene in olfactory organ, gills, skin, spleen and head kidney from 7 days Ich-infected trout. (E) The relative expression of Ich-18SrRNA gene in olfactory organ at 1, 7, 14, 21, 28 and 75 days post infection. Data in d and e are representative of at least three independent experiments (mean and s.e.m.). Statistical analysis was performed by unpaired Student’s *t*-test. \**P* < 0.05, \*\**P* < 0.01 and \*\*\**P* < 0.001.

**S3 Fig.**
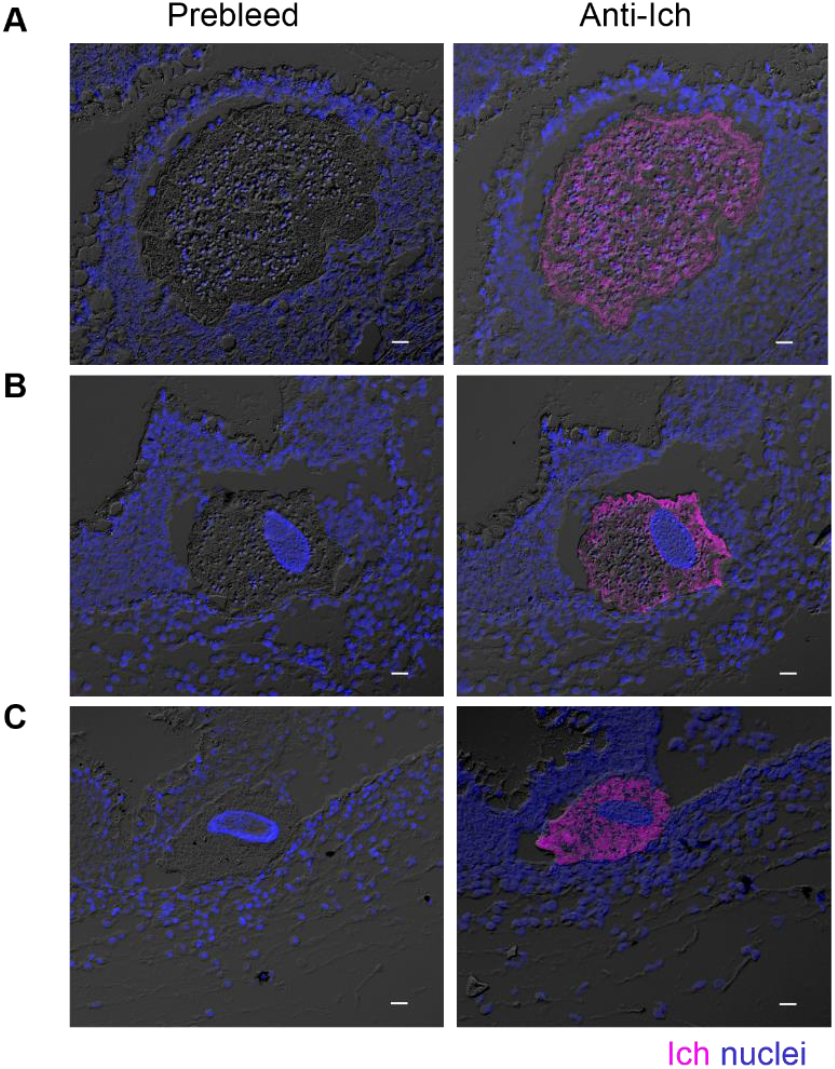
Isotype control staining for anti-Ich antibodies in trout olfactory torgan paraffin sections. Three different microscope images of consecutive slides of prebleed (A-C left) and anti-Ich (A-C right) antibodies staining of Ich parasite in olfactory organ paraffin sections from 28 days Ich-infected fish (*n* = 4). Nuclei were stained with DAPI (blue) and Ich with anti-Ich pAb (magenta). Scale bars, 20 μm. Data are representative of three independent experiments.

**S4 Fig.**
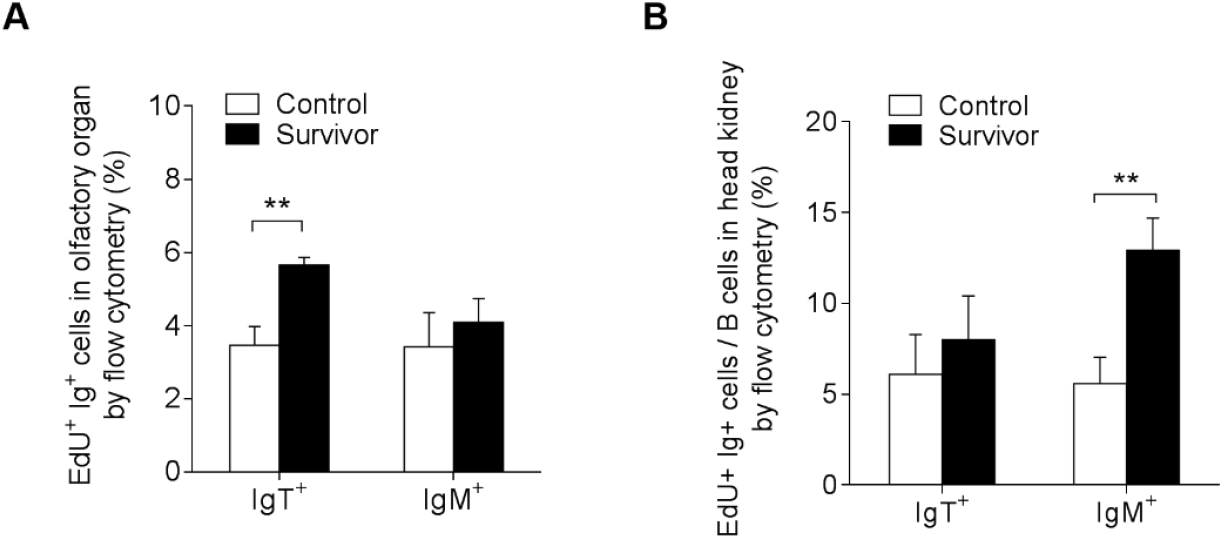
Proliferative responses of IgT^+^ and IgM^+^ B cells in the olfactory organ and head kidney of survivor trout. (A and B) Percentage of EdU^+^ cells from total olfactory organ and head kidney IgT^+^ and IgM^+^ B cell populations in control and survivor fish by flow cytometry analysis (*n* = 9). Data are representative of at least three different independent experiments (mean and s.e.m). Statistical analysis was performed by unpaired Student’s ŕ-test. \**P* < 0.05, \*\**P* < 0.01 and \*\*\**P* < 0.001.

**S1 Table.**
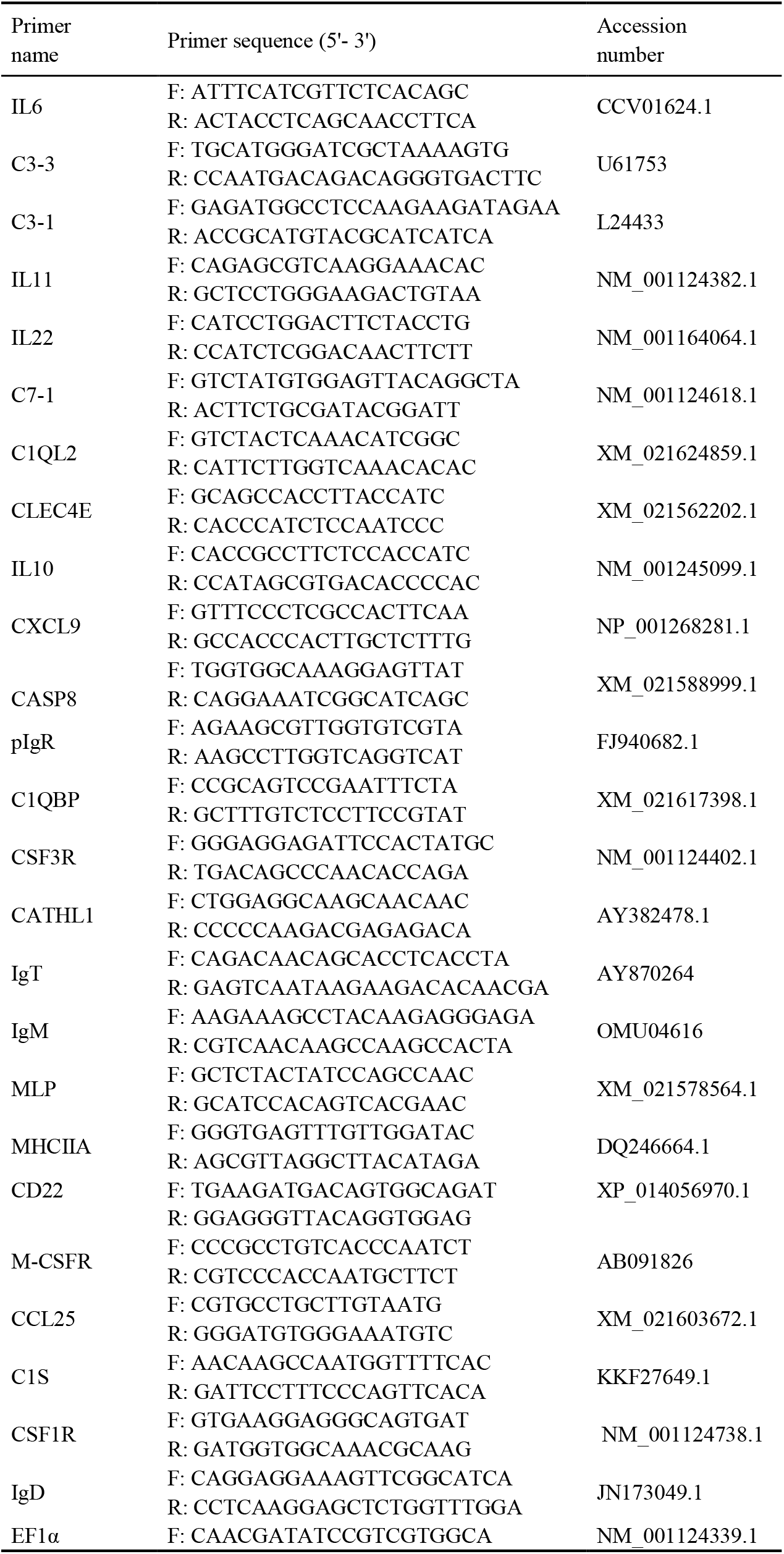
List of primers for real-time quantitative PCR amplifications.

